# Cumulative pregnancy and postnatal environmental exposures impact social behaviour in male mice associated with epigenetic, ribosomal, and immune dysregulation

**DOI:** 10.1101/2025.03.19.644073

**Authors:** Morgan C Bucknor, Brooke A Keating, Velda X Han, Brian S Gloss, Pinki Dey, Nader Aryamanesh, Lee L Marshall, Mark E Graham, Ruwani Dissanayake, Xianzhong Lau, Shrujna Patel, Stela P Petkova, Anand Gururajan, Russell C Dale, Markus J. Hofer

**Author notes:** **Joint senior author** (Russell C Dale, Markus J. Hofer). **Corresponding author:** A/Prof Markus J Hofer.

## Abstract

Environmental exposures across critical developmental windows can significantly influence brain development and contribute to the pathogenesis of neurodevelopmental disorders (NDDs). Emerging clinical evidence suggests that cumulative environmental factors during early development result in more pronounced disease phenotypes in offspring. Here, we developed a ‘triple-hit’ model using C57Bl/6JAusb mice to examine the cumulative effects of antenatal social stress, antenatal chronic high-fat diet consumption, and postnatal poly(I:C) exposure on neurodevelopmental outcomes in offspring. Male ‘triple-hit’ offspring displayed autism-like social deficits and an overall increased susceptibility to neurodevelopmental behavioural alterations in adulthood compared to male non-stressed controls. Conversely, these behavioural changes were not observed amongst female ‘triple-hit’ offspring. Single-cell RNA (scRNA) transcriptomic and bulk proteomic sequencing were performed in male ‘triple-hit’ offspring across peripheral blood immune cells and brain tissue. scRNA sequencing in microglia, astrocytes, and oligodendrocytes revealed dysregulation in critical glial cell processes, ribosomal functions, and chromatin remodelling. Similar functional themes were observed across peripheral blood macrophages, neutrophils, and naïve B cells - displaying immune and ribosomal dysregulation at the transcriptional level. Proteomics pathway enrichment analyses revealed significantly reduced protein abundance in ribosomal biogenesis, translation, and chromatin remodelling functions in both peripheral blood immune cells and brain tissue. We also observed synaptic dysfunction in the blood and brain proteome. Overall, using our unique ‘triple-hit’ model, we demonstrate that multiple early life environmental exposures drive NDD-associated behaviours in a sex-specific manner; and is associated with overlapping central and peripheral molecular mechanisms that have clinical relevance to NDD pathogenesis.

## 1. Introduction

The regulatory processes that govern foetal development are highly orchestrated which render them sensitive to environmental perturbations (Han et al., 2021a; Kalish et al., 2021). An inflammatory intrauterine environment can be triggered by a diverse host of maternal exposures including obesity, gestational diabetes, psychosocial stress, autoimmunity, pollutant exposures or infections during pregnancy (Bronson and Bale, 2016; Chen et al., 2020; Fernandes et al., 2021; Han et al., 2021a). This maternal state, referred to as maternal immune activation (MIA), is hypothesised to disrupt developmental programming of foetal immune and brain cells by priming them towards a more disease susceptible phenotype, thereby increasing an individuals’ lifetime risk of developing neurodevelopmental disorders (NDDs) (Bergdolt and Dunaevsky, 2019; Knuesel et al., 2014; Weber-Stadlbauer, 2017).

NDDs are a lifelong, complex group of learning and behavioural disorders whose multifactorial aetiology is associated with genetic predisposition, the intrauterine experience, and postnatal stressors such as psychological trauma or infection (Bergdolt and Dunaevsky, 2019; Carlsson et al., 2021; Giovanoli et al., 2013; Han et al., 2021b; Toth, 2015). The relative contributions of each factor that translate to overt behavioural impairments has been challenging to ascertain, and available therapies mitigate symptoms but do not target disease mechanisms. Autism spectrum disorder (ASD), attention deficit/hyperactivity disorder (ADHD), tic disorders and Tourette syndrome (TS) are common examples of NDDs and many share overlapping symptomology (Carlsson et al., 2021; Kern et al., 2015; Morris-Rosendahl and Crocq, 2020). Primary core and associated symptoms include impairments in attention, socialisation, communication and anxiety, plus repetitive or ritualistic patterns of behaviour such as counting, repeating and arranging (Frye, 2018; Grzadzinski et al., 2011; Kern et al., 2015). There is also a predominant male sex bias among NDDs; ASD and ADHD for example, are more commonly diagnosed in males than females with sex ratios of 3:1 and 2:1, respectively (Breach and Lenz, 2022).

Peripheral-central nervous system (CNS) immune interactions are imperative for ensuring normal brain development (Han et al., 2021a; Haruwaka et al., 2019; Lehmann et al., 2018; Wohleb and Delpech, 2017; Zengeler and Lukens, 2021). Thus, the assessment of pathological mechanisms between both axes is crucial for understanding NDD pathophysiology. Extensive evidence implicate microglia, the brain’s resident immune cells, as highly susceptible to developmental epigenetic priming in response to MIA, due to their early embryonic colonisation and substantial cellular plasticity (Bilbo et al., 2018; Bordeleau et al., 2020; Knuesel et al., 2014; Perry and Holmes, 2014; Picard et al., 2021; Reemst et al., 2016, 2022; Suzuki et al., 2013a; Thion and Garel, 2017). Neurodevelopmental impacts of other glial cells (i.e., astrocytes, oligodendrocytes) remain largely understudied. Moreover, peripheral immune priming is suggested to co-occur with microglial priming during foetal development representing another crucial component of NDD pathogenesis (Bergdolt and Dunaevsky, 2019; Bordeleau et al., 2020; Estes and McAllister, 2015; Labouesse et al., 2015). Peripheral immune dysregulation is common in NDD patients (Estes and McAllister, 2015; Ju et al., 2025), and with limited access to brain tissue in clinical settings, peripheral blood presents a compelling candidate as a surrogate biomarker for brain pathology.

In this study, we explored the cumulative impacts of multiple environmental factors on long-term neurodevelopmental outcomes in offspring. To achieve this, we developed a novel ‘triple-hit’ MIA model integrating established NDD risk factors, including maternal psychosocial stress before conception, antenatal high-fat (HF) diet exposure, and postnatal immune challenge (poly(I:C), PIC), to better reflect the human experience (Bucknor et al., 2024; Han et al., 2021b; Nielsen et al., 2024). In addition to behavioural phenotyping, we characterised changes to the omics landscape in male peripheral blood and brain cells to capture disease promoting peripheral-CNS mechanisms. Phenotypic analysis revealed that ‘triple-hit’ MIA male offspring were more susceptible to developing NDD-associated behavioural deficits compared to non-stressed controls and their female counterparts. Furthermore, single-cell RNA transcriptomics and proteomic sequencing revealed alterations to peripheral immune functions, glial cell dysfunction, dysregulated ribosomal processes, synaptic homeostasis and chromatin remodelling in both the brain and blood of these offspring; all relevant pathways that share significant associations with the pathogenesis of various NDDs (Chen et al., 2019; Han et al., 2021a; Kundakovic and Jaric, 2017; Meltzer and Van De Water, 2017; Mossink et al., 2021). Altogether, these findings emphasise the systemic impacts of early life adversity and offers novel insights into the potential of peripheral blood to serve as a reliable guide for future NDD therapeutic interventions.

## 2. Materials and methods

### 2.1 Animals and ethical approval

All animals were housed in a specific pathogen-free (SPF) environment at the Charles Perkins Centre (Sydney, Australia) animal facility (20-24℃, 40-70% humidity, 12h light:dark cycle). All male and female offspring in this study were generated via on-site breeding from C57Bl/6Ausb dams as previously reported (Bucknor et al., 2024). Male and female mice used for breeding were obtained from Australian BioResources (ABR; Moss Vale, NSW). All experiments were performed in compliance with the NSW Animal Research Act and the 8^th^ edition of the NHMRC ‘Australian Code of Practice for the care and use of animals for scientific purposes.’ This study was reviewed and approved by the University of Sydney Animal Ethics Committee.

### 2.2 Maternal and offspring diets

Dams had *ad libitum* access to either a high-fat, no soy semi-pure diet (HF+ diet, 43% kcal fat diet, SF21-208, Specialty Feeds, WA, Australia) or a low-fat, high-sugar, no soy semi-pure control diet formulation (HF-diet, 12.3% kcal fat diet, SF21-209, Specialty Feeds, WA, Australia) prior to, during gestation and during the lactation period. The detailed composition of maternal diets and time of exposure has been previously described (Bucknor et al., 2024). F1 generation offspring were weaned at postnatal day (PND) 28 and transitioned to standard, grain-based laboratory chow (SF00-100, Specialty Feeds, WA Australia) for the remainder of the study.

### 2.3 Offspring pre- and postnatal stress

In brief, dams were randomised to either chronic high-fat diet consumption (HF+ or HF-), six-weeks of social instability stress (SIS+ or SIS-) or both in parallel as described previously (Bucknor et al., 2024). This generated four distinct maternal stress groups: high-fat diet fed only (HF+/SIS-), high-fat diet fed and social instability stressed (HF+/SIS+), low-fat diet fed only (HF-/SIS-), and low-fat diet fed and social instability stressed (HF-/SIS+). At ≥16 weeks of age, females (n = 45) were paired for breeding and left undisturbed until birth of litters. Following birth of a litter, offspring were left undisturbed until weaning. The day after weaning (PND 29), offspring were randomised to receive either a single intraperitoneal injection of 10 mg/kg body weight poly(I:C) (high molecular weight, Cat# tlrl-pic-5, InvivoGen) or endotoxin-free 0.9% NaCl (Baxter, Australia). Following poly(I:C) or vehicle exposure, adult offspring were behaviourally phenotyped and then euthanised for tissue collection. Further methodological details describing the maternal prenatal stress model, and poly(I:C) administration can be obtained in **Extended Methods**. Offspring were group-housed with littermates and/or same-sex treatment group cage mates from other litters where possible for the remainder of the study. Refer to study timeline (**Fig. 1**) for experimental design and **Table 1** for sample group information.

**Figure 1.**
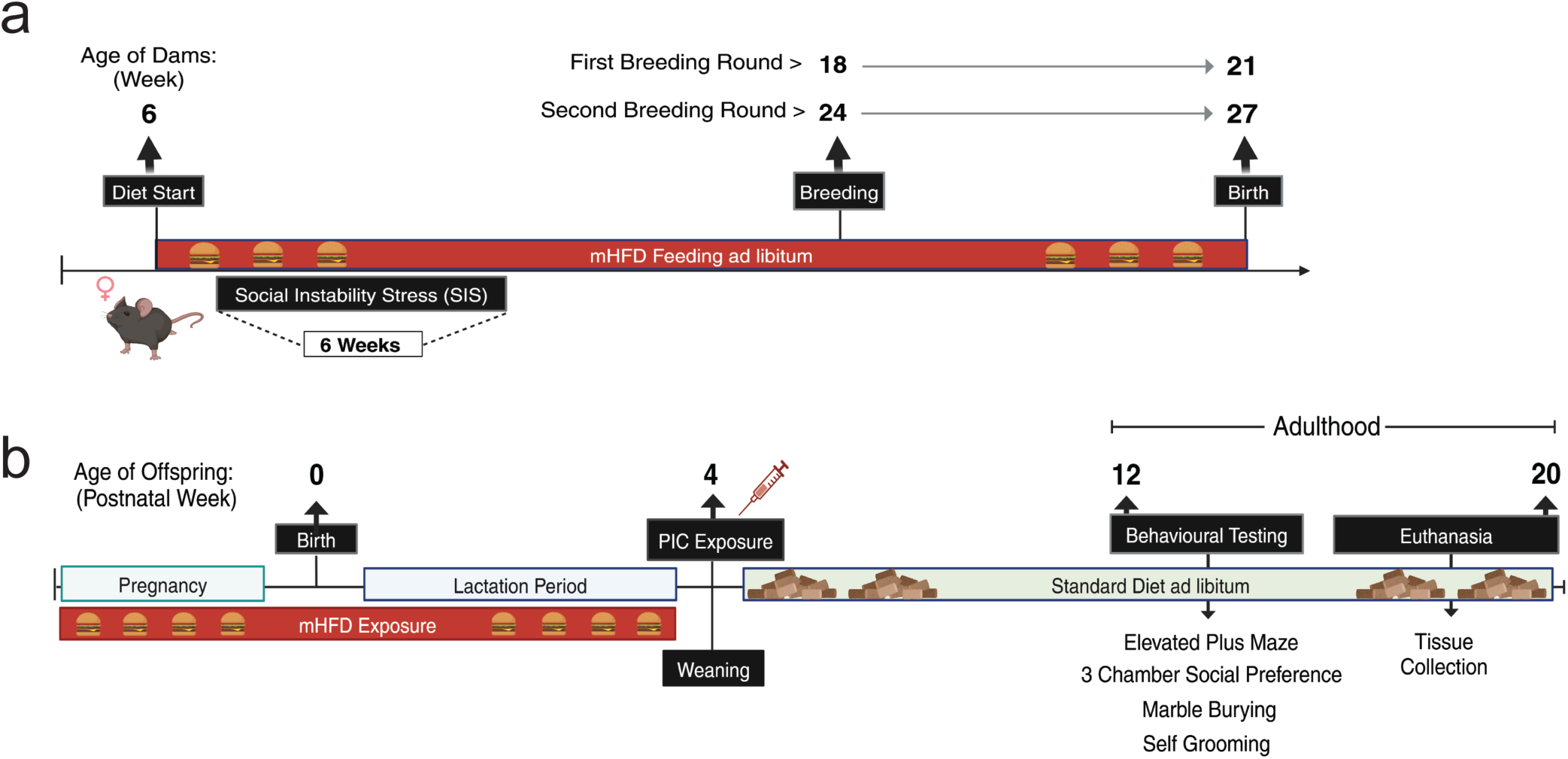
Experimental timeline. (**a**) Maternal high-fat diet and social instability stress paradigm to generate HF+/SIS+ and/or HF+/SIS- females. Dietary consumption began at 6 weeks of age and continued into the lactation period. SIS exposure lasted for 6 weeks prior to inducing pregnancy. HF+ females were staggered for breeding 2 weeks apart, the first round of matings began at 18 weeks of age. Second round of matings paired females at 24-weeks of age. (**b**) Neonates were exposed to mHFD via maternal milk and left undisturbed until weaning. At 4 weeks of age offspring were randomised to receive 10mg/kg of poly(I:C) or NaCl vehicle solution via intraperitoneal injection. Behavioural testing began at 12 weeks of age, followed by euthanasia for tissue collection. Created with biorender.com.

**Table 1.** Developmental window of environmental exposures, sample group information and number of offspring.

| Preconception | Antenatal | Postnatal | ♂<sup>c</sup> | ♀<sup>d</sup> |
| --- | --- | --- | --- | --- |
| mSIS<sup>a</sup> | mHFD<sup>b</sup> | Poly(I:C) | (n) | (n) |
| - | - | - | 5 | 7 |
| + | - | - | 12 | 12 |
| - | + | - | 5 | 1 |
| - | - | + | 8 | 10 |
| + | + | + | 5 | 6 |
|  |  |  | N= 35 | N = 36 |

<sup>a</sup>mSIS = maternal social instability stress.
<sup>b</sup>mHFD = maternal high-fat diet.
<sup>c</sup>♂ = male.
<sup>d</sup>♀ = female.

### 2.4 Behavioural testing

Behavioural testing began on both male and female offspring at 12-weeks of age and was carried out over a 2-week period. The same researcher conducted all tests and was blinded to offspring group conditions. All tests were performed during the light cycle between 0900 and 1700h. To assess NDD-associated behaviours, we selected a battery of tests relevant to the symptomatology observed in NDDs (Silverman et al., 2010): elevated plus maze (anxiety-like behaviour), 3-chamber social preference (social engagement), marble burying and self-grooming (repetitive behaviours), with 2 - 4 days of rest between tests. Detailed description of all assay procedures is provided in brief below and expanded in **Extended Methods**. Raw data of elevated plus maze and 3-chamber social preference test was analysed using AnyMaze video tracking software (V7.20). Time spent grooming was manually scored using Behavioural Observation Research Interactive Software (Boris). The number of marbles buried was recorded by two independent researchers at the end of the testing period.

### 2.4 Elevated plus maze test

Elevated plus maze testing was performed as described previously (Bucknor et al., 2024). Briefly, this test allows for assessment of anxiety-like conflict behaviour by allowing the mice to choose between entering the two open arms of the maze (natural exploratory drive) or entering and remaining in the safety of the two closed arms (Flannery et al., 2015). The parameters measured included total distance travelled, percentage of open arm entries and percentage of time spent in the open arms of the maze.

### 2.5 3-chamber social preference

Social impairments are one of the core symptoms of ASD and rodents regularly engage in high levels of social reciprocity (Silverman et al., 2010; Xuan and Hampson, 2014). The 3-chamber social preference test assays for changes in sociability (time spent engaging with a novel mouse rather than an inanimate object) and social novelty (time spent engaging with a novel mouse rather than a familiar mouse) using a 3-chamber apparatus.

This test includes three sequential 10-minute trials beginning with habituation, sociability and lastly, social novelty. The test mouse is placed in the centre chamber and allowed to freely explore both side chambers during the test. The habituation trial allows the mouse to freely explore the apparatus, with each side chamber consisting of only an empty wire cup. Following the habituation trial, the test mouse is confined to the centre chamber while a novel object (golf ball) is placed under one wire cup and a novel sex-matched mouse is placed under the other wire cup in the other side chamber. This trial assesses changes in sociability by measuring time spent in the chamber containing the novel object and novel mouse. For the last trial, the test mouse is confined to the centre chamber while the novel object is replaced with another novel sex-matched mouse. This trial assesses changes in social novelty or recognition by measuring the time spent in the chamber containing the unfamiliar mouse and familiar mouse.

### 2.6 Marble burying

Repetitive or ritualistic behaviours are clinically relevant symptoms of NDDs (Silverman et al., 2010). Therefore, to evaluate repetitive digging behaviours, we performed the marble burying test. The test mouse was placed in a standard cage with bedding 3-4 cm thick and overlaid with 20 black, glass marbles in a 4×5 arrangement. The number of marbles buried by test mice (defined as more than 2/3rds or more covered by the bedding) at the end of the 30 minute test period was recorded.

### 2.8 Self-grooming

Spontaneous, unusually long repetitive bouts of grooming were scored over a period of 20 minutes in a novel, empty cage under dim lighting (< 50 lux) to assess repetitive, obsessive-compulsive like behaviour for each test mouse. The first 10 minute period was unscored and considered the habituation phase, while the remaining 10 minutes were scored.

### 2.9 Integrated neurodevelopmental disorder (NDD) behavioural index

Integrated behavioural measures from complimentary tests can be z-normalised to reduce behavioural noise that is a common artifact of behavioural assays (Guilloux et al., 2011; Kraeuter, 2023). This approach provides an overall measure of the animals’ emotionality dependent on the context of the behavioural tests integrated - which here includes anxiety-like behaviour (elevated plus maze), repetitive behaviours (marble burying and grooming), and social engagement (3-chamber social preference). By integrating these NDD-associated behavioural outputs, we provide an index that summarises each groups overall behaviour relative to controls. Refer to **Extended Methods** for formula used.

### 2.10 Tissue collection for single-cell RNA sequencing and bulk proteomics

For single cell RNA (scRNA)-seq and bulk proteomic analyses, peripheral blood leukocytes and total forebrain tissue were analysed from male offspring for HF+/SIS+/PIC+ and HF-/SIS-/PIC- for two reasons: (i) the male-biased prevalence of NDDs and (ii) it was evident from our behavioural analyses that the cumulative effects of pre- and postnatal stress had less impact on female offspring. scRNA-sequencing was performed using the HIVE™ Single Cell platform (Honeycomb Biotechnologies, Inc. USA). Bulk LC-MS/MS proteomics leveraged an untargeted approach and used the Dionex UltiMate 3000 HPLC system. Complete sample preparation information for scRNA-sequencing and proteomics is provided in **Extended Methods**.

For tissue collection, mice were deeply anesthetised with isoflurane (5% induction, 2.5% maintenance) (IsoFlo®, Abbott Laboratories, Botany, NSW, Australia). Up to 1 mL of peripheral whole blood was collected via cardiac puncture and placed into a 1.5 mL collection tube containing anticoagulant (10% blood volume) (Heparin, Sigma, H3393) on ice. The mouse was then perfused with PBS and the whole brain was dissected. The cerebellum and olfactory bulbs were excised along the anterior to posterior axis from approximately +1.50 mm to −3.50 mm, leaving total forebrain tissue. Regions left intact included cerebral cortices, thalamus, hypothalamus, striatum and midbrain. The tissue was weighed and recorded before tissue homogenisation (gentleMACS Octo Dissociator with Heaters, Miltenyi Biotec, 130-096-427). A single-cell suspension was prepared using the Adult Brain Dissociation kit as per manufacturer’s instructions (Miltenyi Biotec, 130-107-677). To isolate peripheral leukocytes from whole blood, red blood cell lysis was performed using ammonium chloride solution (StemCell Technologies, 07850) at a volume:volume ratio of 9:1. The solution incubated at 4°C for 10-minutes, then cells were washed in Dulbecco’s phosphate-buffered saline containing Ca^2+^, Mg^2+^, glucose and pyruvate (DPBS, ThermoFisher, 14287080) by centrifugation at 250 x g for 6-minutes at 4°C.

### 2.11 scRNA-sequencing: HIVE CLX™ sample capture, library preparation and sequencing

Once single-cell suspensions were generated for each sample condition, samples were loaded into the HIVE collectors according to the manufacturer’s instructions. Approximately 30,000 cells were loaded directly into a designated HIVE collector (Honeycomb Biotechnologies, Inc. USA) in 1% FBS in DPBS. Collectors were centrifuged at 30xG for 3 minutes to allow single-cells to settle into picowells of the HIVE collector, which contained mRNA-capture beads. Media was removed from the collector and 2 mL of CLX sample wash solution (provided by manufacturer) was added. After removing the wash solution, 1 mL of cell preservation solution (provided by manufacturer) was added (<u>Honeycomb Biotechnologies, Inc. USA_Sample Capture protocol</u>). HIVE collectors were stored at −80°C until transferred to the Australian Genome Research Facility Ltd. (AGRF, Westmead, Australia) for transcriptome recovery and the library preparation.

All HIVE CLX^TM^ devices were processed according to the manufacturer’s instructions (<u>Honeycomb Biotechnologies, Inc. USA: Transcriptome Recovery and Library</u> <u>Preparation protocol)</u>. Briefly, HIVE collectors were sealed with a semi-permeable membrane, allowing for the addition of strong lysis solution and hybridization solution. Capture beads with transcripts were extracted from the collector by centrifugation. The remaining library preparation steps were performed in a 96-well format. The size profiles of the final libraries were determined on a TapeStation platform with a D5000 ScreenTape System (Agilent Technologies, Santa Clara, CA, USA), and the concentration of the final pooled library was determined by qPCR assay before sequencing on the Illumina NovaSeq X sequencing platform with custom primers (provided in the HIVE single-cell RNA seq processing kit) (AGRF, Melbourne, Australia).

### 2.12 scRNA-sequencing bioinformatics analysis

Raw base calls for brain and blood samples were processed using the Illumina BCL Convert 4.1.5 pipeline to generate FastQ files. FastQ files were analysed using BeeNet (v1.1.3) software, mapped to the reference *Mus musculus* genome (mm10.104) and imported into the R statistical environment (v4.3.1) using the *Seurat, tidyverse* and *patchwork* R packages. Preprocessing was performed according to beenet v1.3 workflow. Briefly, before expression data was merged, cells with a high mitochondrial transcript ratio (> 0.20), less than 300 genes or 600 transcripts were excluded. Merged data were normalized and scaled using *SCtransform* before clustering and UMAP projection. ScType was used to assign cell types using the *auto_detect_tissue_type* workflow (Ianevski et al., 2022). Data was split by cell type and then renormalized, markers between conditions were determined using *FindMarkers* with the parameters logfc.threshold = 0,test.use = "wilcox",min.pct = 0.05. Pathway enrichment was determined via over representation analysis (ORA) to obtain Gene Ontology (GO) pathways for differentially expressed genes (adj p value <0.05) using *enrichGO* in the *CompareCluster* function from ClusterProfiler (Wu et al., 2021).

### 2.13 Protein lysate sample preparation for proteomics

Following brain and peripheral blood leukocyte cell isolation as described above, the remaining fraction of cells were lysed for protein for unbiased mass spectrometry analysis. Cell suspensions were incubated in 200 μL of 1X Lysis Buffer containing: 0.8% v/v Triton X-100, 50 mM HEPES (pH 7.4) with NaOH, EDTA free protease inhibitor, PhosSTOP and 2 μL of 2 mM phenylmethyl sulfonyl fluoride in ethanol (PMSF) at 85°C for 10 minutes on a heating block. Samples were then stored at −80°C until LC-MS analysis. Extended details are provided in **Extended Methods** concerning tissue lysis, protein digestion and peptide tagging for LC-MS/MS analysis.

### 2.14 LC-MS/MS analysis

LC-MS/MS was performed using a Dionex UltiMate 3000 RSLC nano system and Q Exactive Plus hybrid quadrupole-orbitrap mass spectrometer (ThermoFisher Scientific). Each HILIC fraction was loaded directly onto an in-house 300 mm long 0.075 mm inside diameter column packed with ReproSil Pur C18 AQ 1.9 μm resin (Dr Maisch, Germany). The column was heated to 50°C using a column oven (PRSO-V1, Sonation lab solutions, Germany) integrated with the nano flex ion source with an electrospray operating at 2.3 kV. The S lens radio frequency level was 50 and capillary temperature was 250°C.

All brain and blood sample fractions were analysed using data-dependent acquisition LC-MS/MS. MS scans were performed at 70,000 resolution with an automatic gain control target of 1,000,000 for a maximum ion time of 100 ms (*m*/*z* 375 to 1500). MS/MS scans were at 35,000 resolution with an automatic gain control target of 200,000 and maximum ion time of 100 ms. The loop count was 12, the isolation window was 1.1 m/z first mass was fixed at 120 m/z, and the normalized collision energy was 31. Singly charged ions and those >8+ were excluded, with a 35 s dynamic exclusion.

### 2.15 Differential abundance protein and pathway enrichment analysis

For data cleaning, the raw ‘proteinGroups.txt’ LC-MS/MS output file was processed from MaxQuant v1.6.7.0. Each protein group must have had at least one unique peptide. Proteins were removed from further analysis if they matched contaminant or reverse sequence decoy entry prefixes (CON_ or REV_). Proteins with one or more missing values in any samples were also removed.

Data was normalised by first identifying the UniProt accession for each protein group following rules from Engholm-Keller et al (2019) (Engholm-Keller et al., 2019). Multi-mapped proteins were also excluded to avoid false enrichment due to alignment errors. The samples were first log (base 2) transformed, then between sample normalisation was performed using the ‘scaled’ normalisation from the *limma* R package (v4.2.3). To remove batch effects from biological data, the remove unwanted variation *RUV* R package was used (Gagnon-Bartsch and Speed, 2012).

Differential abundance analysis of proteins was performed using the adjusted abundance matrix. The results from this differential abundance analysis were then used for all analyses. For linear modelling, differential abundance analysis of proteins was performed using the *limma* R package. The linear model for comparing each pair of time points was fitted using the *lmFit* function and p-values calculated using the empirical Bayes method using the ‘eBayes’ function. Significant differentially expressed proteins were defined by (q-value <0.05). Differentially expressed proteins were analysed for GO pathway enrichment analysis using the *enrichGO* function from the *clusterProfiler* R package.

### 2.16 Statistical Analysis

No statistical methods were used to predetermine offspring sample sizes as this was limited by breeding outcomes from dams described previously (Bucknor et al., 2024). All statistical testing was performed in the R statistical environment (v2023.12.1+402). GraphPad Prism software (v10.4.0) was used for graphical representation of physiological and behavioural phenotyping results. For physiological phenotyping, our analyses employed a mixed-effects model using the *lme4* and *car* R Package to analyse the effect of age, group and their interaction on weight gained. The subject (mouse) was included as a random effect. Behavioural output from the elevated plus maze, marble burying, self-grooming and integrated z-score met assumptions of normality (Shapiro-Wilk test) and homogeneity of variance (Levene’s test) – therefore, one-way ANOVA was performed. Where a significant main effect of group was detected, Dunnett’s post-hoc test was performed. For social preference testing, we first fit a linear mixed-effects model using the (*lme4*) R Package and detected zero variance when including the random effect of each mouse. Therefore, a linear model (lm) was adopted which focused on fixed effects only (group and social zone). We then conducted Type III ANOVA on the model output to assess main effects (group) and interactions (group x social zone). Where significant main effects or interactions were detected, Tukey’s post-hoc test was performed.

## 3. Results

### 3.1 Early life exposure to mHFD or poly(I:C) alone leads to increased time spent in the open arms of the plus maze

Mice underwent behavioural testing using NDD-associated assays at 12-weeks of age, starting with the elevated plus maze (Silverman et al., 2010). Description of male behaviours are provided below, while complementary results from female offspring are provided in supplementary material **(Suppl. Fig. S3-4)**. Anxiety is not considered a core symptom of NDDs but is frequently reported as an associated symptom, depending on diagnostic criteria (Kern et al., 2015). When we performed elevated plus maze testing, we found no statistically significant differences in the percentage of open arm entries between groups (**Fig. 2a**). However, one-way ANOVA revealed a main effect of group on percentage of time spent in the open arms (F(4, 30) = 2.913, p = 0.0379). Dunnett’s post-hoc test revealed HF-/SIS-/PIC+ and HF+/SIS-/PIC- offspring spent more time in the open arms relative to HF-/SIS-/PIC- controls (14.9% (p = 0.030) and 17.6% (p = 0.020) mean increase in time, respectively (**Fig. 2b**).

**Figure 2.**
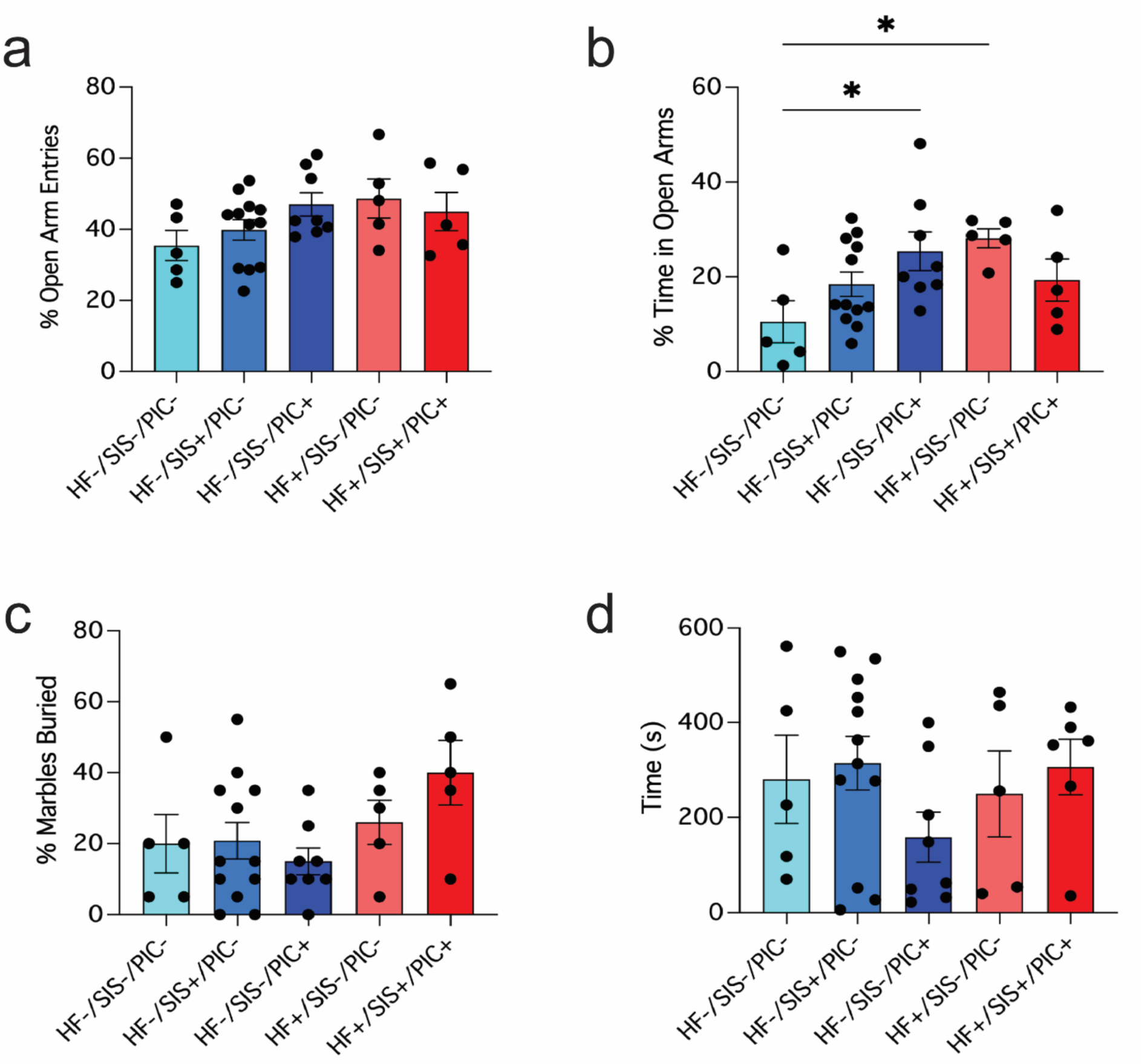
Male offspring exposed to poly(I:C) or mHFD in isolation of other factors display increased time spent in open arms of plus maze. (**a-b**) Elevated plus maze, (**a**) percentage of open arm entries and (**b**) percentage of time spent in the open arms, (*p<0.05); (**c**) marble burying test – percentage of marbles buried; (**d**) self-grooming test - time(s) spent grooming. (**a**-**d**) One-way ANOVA followed by Dunnett’s post-hoc test. Data shown as mean ± SEM; each individual data point represents one mouse.

Repetitive behaviours were assayed using the marble burying test and observing grooming behaviours. Here, the marble burying test did not reveal any statistically significant differences between groups and nor did time spent grooming differ significantly (**Fig. 2c-d**). Overall, based on these tests, the most significant alterations in behaviour related to increases in exploratory drive in offspring exposed to only poly(I:C) or mHFD.

### 3.2 HF+/SIS+/PIC+ offspring demonstrate deficits in social behaviours

A linear model was used to examine the effects of group and zone on time spent either with a novel mouse or novel inanimate object during the sociability trial. Type III ANOVA of this model revealed significant main effects of group (F(4, 60) = 4.80, p = 0.0019), zone (F(1, 60) = 22, p = p<0.001) and interaction between group x zone (F(4, 60) = 7.84, p<0.001). Tukey’s post-hoc test revealed significantly more time spent in the novel mouse zone versus novel object zone in HF-/SIS-/PIC- controls (+140.38s (p<0.0001)), HF-/SIS+/PIC- and HF-/SIS-/PIC+ offspring (+109.89s and +158.66s (p<0.001). This profound social engagement with the novel mouse is expected. Whereas, both HF+ groups did not spend significantly more time with the novel mouse, instead they spent almost equal amounts of time in both zones (**Fig. 3a**). These changes in sociability suggest that exposure to mHFD alone, as well as in the context of additional mSIS and poly(I:C), result in social exploration deficits.

**Figure 3.**
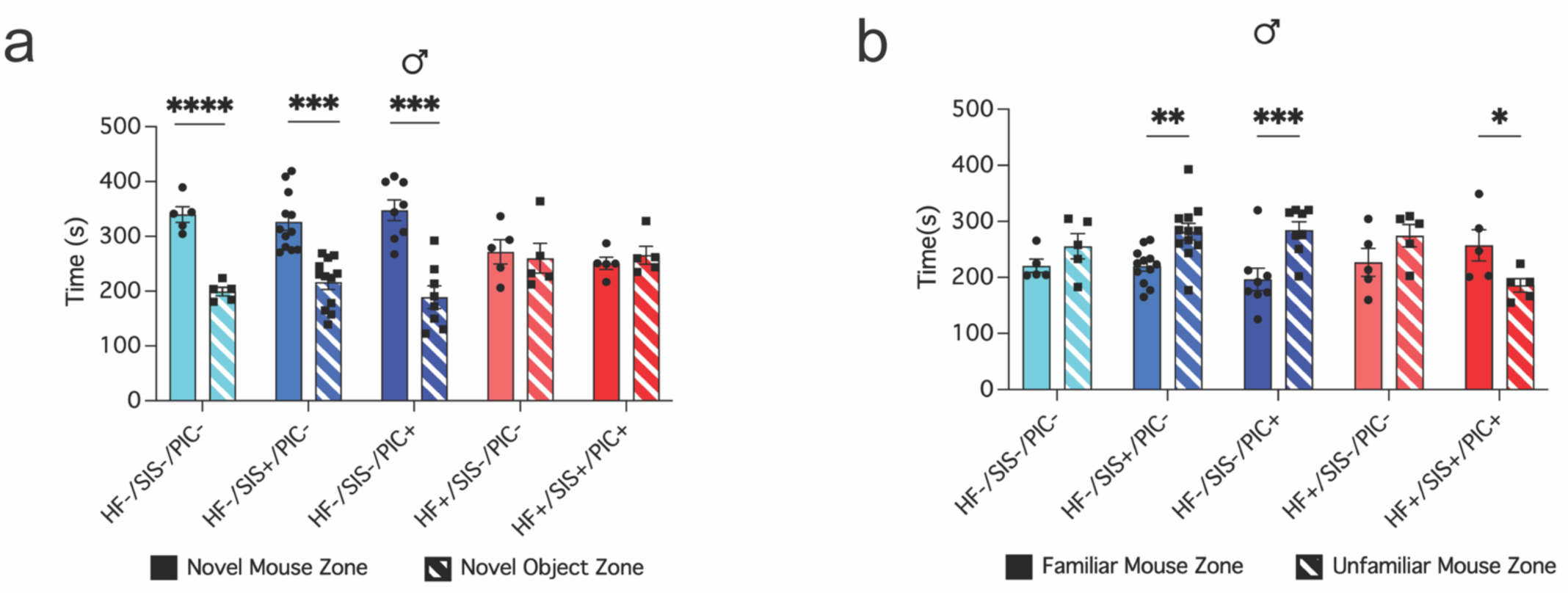
Male HF+/SIS+/PIC+ offspring display deficits in social recognition and sociability. (**a-b**) 3-chamber social preference test, (**a**) total time (s) spent in novel mouse zone and novel object zone within group comparisons, (***p<0.001, ****p<0.0001); (**b**) total time (s) spent in familiar mouse zone and unfamiliar mouse zone within group comparisons, (*p<0.05, **p<0.01, ***p<0.001) (**a-b**) Linear model followed by type III ANOVA and Tukey’s post-hoc test. Data shown as mean ± SEM; each individual data point represents one mouse.

The same statistical approach was used for analysing the effects of group and social zone on time spent either with an unfamiliar or familiar mouse. Type III ANOVA revealed a significant interaction between group x zone (F(4, 60) = 5.139, p = 0.0013). Upon further investigation, Tukey’s post-hoc test revealed HF-/SIS+/PIC- and HF-/SIS-/PIC+ spent significantly more time with the unfamiliar mouse than the familiar mouse [-61.5s (p = 0.0017) and −88s (p = 0.0003)]. This distinction in time spent with familiar versus unfamiliar shows intact recognition and novelty interest. HF+/SIS+/PIC+ offspring, in contrast, spent significantly more time in the familiar mouse zone compared to the novel (71.46s (p = 0.0165). These findings suggest that the cumulative effects of all stressors (HF+/SIS+/PIC+) lead to impairments in social memory or recognition (**Fig. 3b**).

### 3.3 Integrated z-score indicates increased risk of neurodevelopmental disorder– like behaviours in adult male HF+/SIS+/PIC+ offspring

To best interpret our behavioural findings across all tests, we performed an integrated z-score calculation. To do this we z-normalised each individual animal’s behavioural output, added them together and divided by the number of tests included in the integration (see **Extended Methods**). Since our behavioural phenotyping consisted of NDD-associated tests, we described this output as an integrated NDD behavioural index. Values >1 are indicative of increased susceptibility to NDD-like behaviours, < 1 less susceptible, and 0 is no change. When we assessed the difference in z-score across male groups, one-way ANOVA detected a significant main effect of group (F(4, 30) = 5.73, p = 0.0015). Overall, HF+/SIS+/PIC+ males exhibited a statistically significant increase in z-score compared to HF-/SIS-/PIC- controls, and Dunnett’s post-hoc test confirmed this (+1.5 standard deviation difference (p = 0.0101) (**Fig. 4a**). When we performed the same analysis in female counterparts, we found the absence of any increased risk for neurobehavioural impairments. Instead, all female groups clustered closely around zero (**Fig. 4b**). Lastly, to see if sex may influence NDD-like behavioural susceptibility, we fit a linear model followed by type III ANOVA which revealed a significant interaction between sex x group (F(4, 60) = 4.99, p = 0.001). Tukey’s post-hoc test revealed female HF+/SIS+/PIC+ offspring displayed a significantly lower average z-score compared to male HF+/SIS+/PIC+ offspring (−1.4 standard deviation difference (p = 0.0003) (**Fig. 4a-b**). These results indicate firstly, that mHFD, mSIS and postnatal poly(I:C) exposure differentially impact males and females; and male HF+/SIS+/PIC+ offspring are most vulnerable to NDD-associated behavioural impairments compared to HF-/SIS-/PIC- controls.

**Figure 4.**
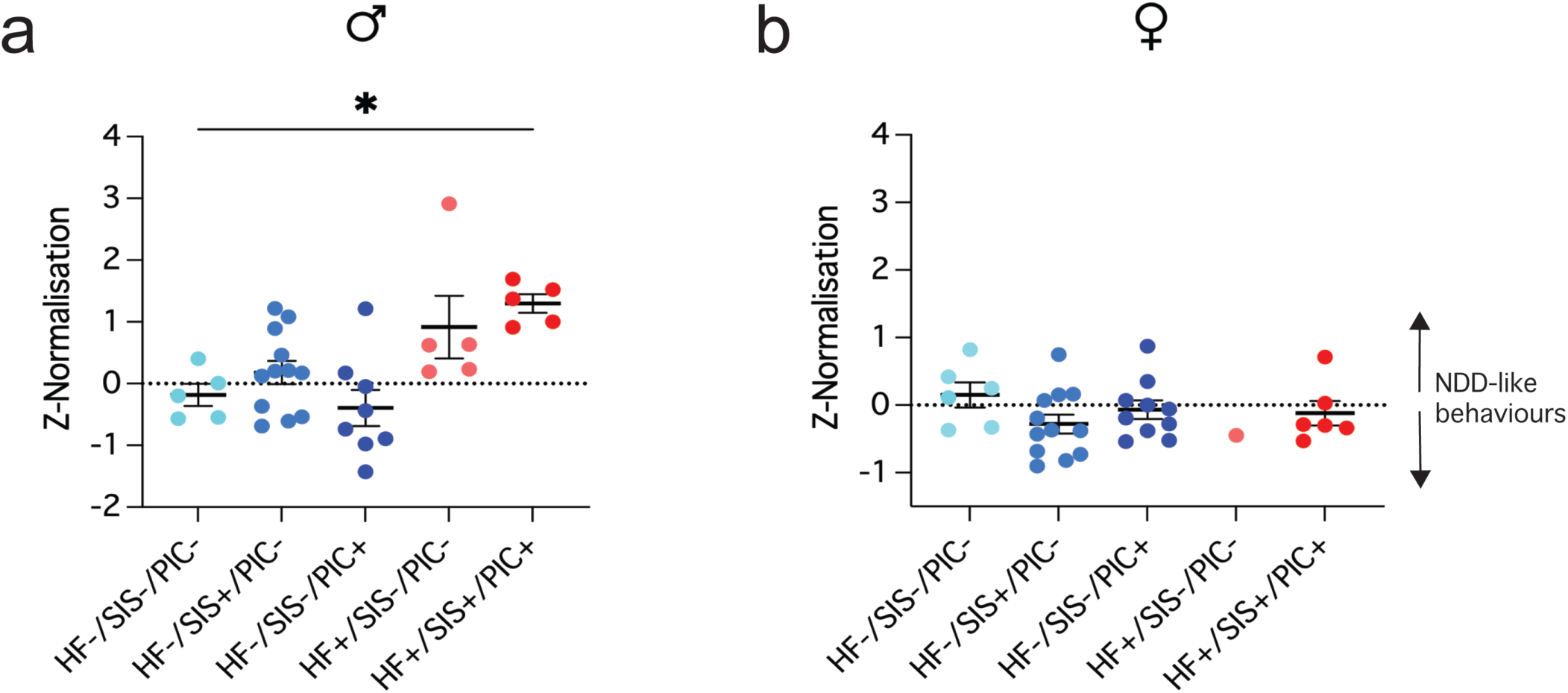
Male HF+/SIS+/PIC+ offspring exhibit increased susceptibility to neurodevelopmental abnormalities compared to HF-/SIS-/PIC- controls, with no similar effect observed in females. (**a**-**b**) Integrated NDD-behavioural z-score incorporating elevated plus maze, marble burying, self-grooming and social preference testing output. (**a**) (*p<0.05); (**b**) no significant difference. (**a**-**b**) One-way ANOVA followed by Dunnett’s post-hoc test. Data shown as mean ± SEM; each individual data point represents one mouse. Males (♂), females (♀).

### 3.4 scRNA sequencing in brain glial cells reveals dysregulation in glial function, mRNA processing and epigenetic regulation pathways

Following our behavioural phenotyping results that revealed adult male HF+/SIS+/PIC+ offspring to be especially vulnerable to NDD-associated behavioural deficits, we sought to investigate what biological pathways in specific cell types could be mediating this phenotype relative to HF-/SIS-/PIC- controls (n = 2/group). To do this, we characterised the transcriptomic profile of the forebrain at single-cell resolution. Relevant quality control information is provided in **(Suppl. Fig. S6a-c).** Further information regarding sample preparation is provided in **Extended Methods.**

Cell clustering and uniform manifold approximation and projection (UMAP) analysis of samples revealed 11 distinct cell type clusters (**Fig. 5a**). Statistically significant (FDR <0.05) differentially expressed genes (DEGs) ranged from 12 to 1333 across captured cell types in HF+/SIS+/PIC+ versus HF-/SIS-/PIC- offspring, with the highest proportions of DEGs found in oligodendrocytes, astrocytes and microglia (**Fig. 5b**). Therefore, we restricted our functional gene ontology (GO) pathway enrichment using Over-representation Analysis (ORA) from these DEGs to these 3 primary glial cells.

**Figure 5.**
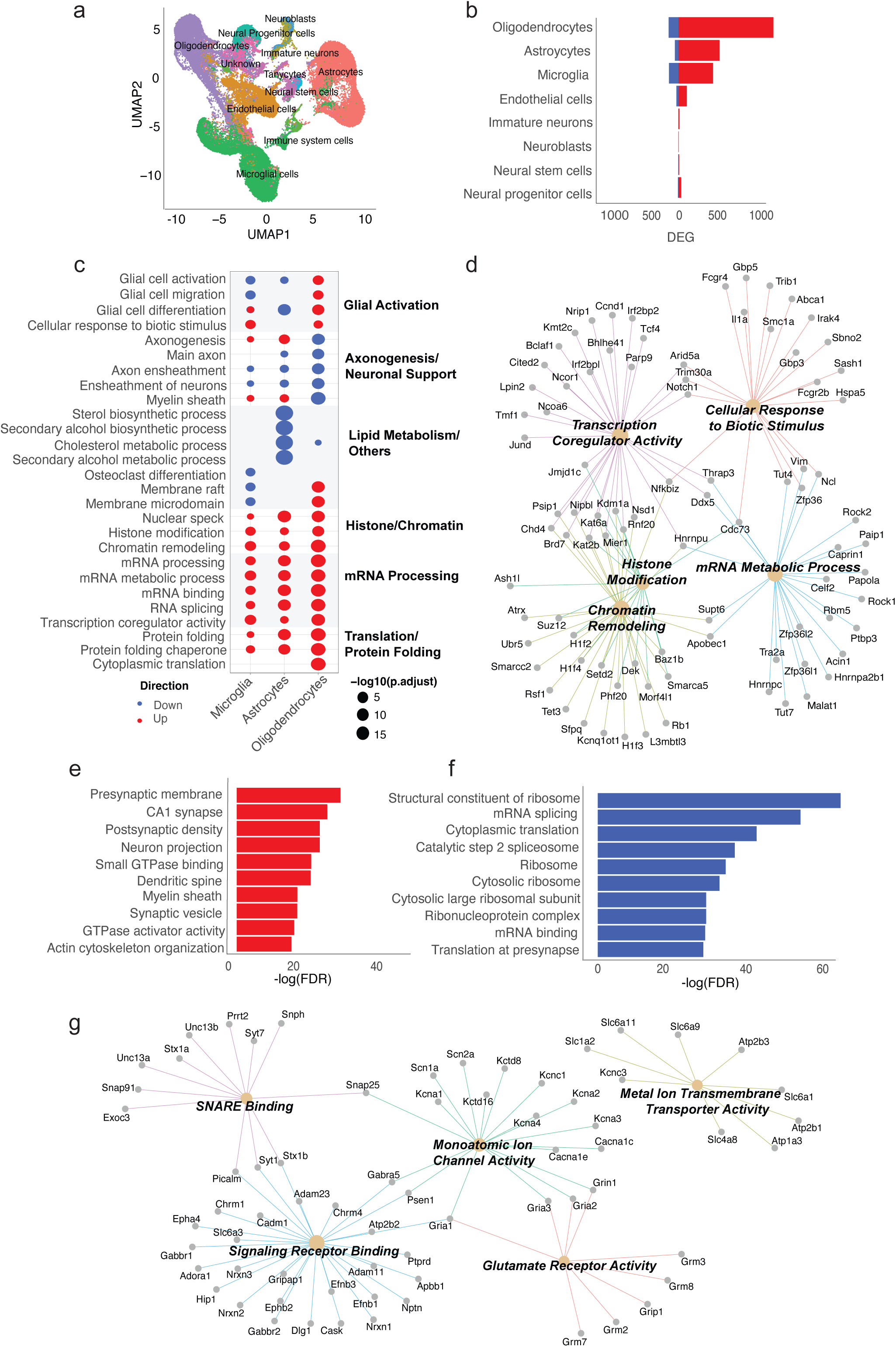
Brain transcriptome and proteome in male HF+/SIS+/PIC+ offspring versus controls. (**a**) UMAP (uniform manifold approximation of projection) of single-cell populations captured for sequencing: astrocytes, endothelial cells, immature neurons, immune system cells, microglial cells, neural progenitor cells, neural stem cells, neuroblasts, oligodendrocytes, tanycytes, unknown (**b**) proportions of significantly differentially expressed genes (DEGs) (FDR <0.05) across brain cell types. Oligodendrocytes (1333), astrocytes (567) and microglia (560) display largest proportions of upregulated DEGs. Colours indicate DEG direction of change (*red*: up; *blue*: down). (**c**) Dot plot displaying GO analysis of DEGs (FDR <0.05) HF+/SIS+/PIC+ versus HF-/SIS-/PIC- controls in select cell types (microglia, astrocytes, oligodendrocytes) presented on x-axis. Enriched pathway terms are displayed on left y-axis and clustered biological themes of like terms presented on right y-axis. Size of each coloured dot is indicative of the negative log10 of enriched pathways adjusted p-value. The colour of each dot is indicative of the direction of change. (**d**) Connectivity network (CNET) plot of top five upregulated GO pathways and corresponding genes that enrich for each pathway in microglia: transcription coregulator activity, cellular response to biotic stimulus, mRNA metabolic process, chromatin remodelling, histone modification. Each enriched pathway is represented by the respective colours and corresponding genes’ adjusted p-value. (**e**) Bar plot of bulk proteome depicting top 10 enriched GO pathways amongst proteins with increased abundance. Statistical significance depicted by negative log(FDR). (**f**) Bar plot of bulk proteome depicting top 10 enriched GO pathways amongst proteins with decreased abundance. Statistical significance depicted by negative log(FDR). (**g**) Subclustered CNET plot of the upregulated GO pathway ‘presynaptic membrane’ in proteome. Subclusters are based on molecular function and include: SNARE binding, signalling receptor binding, glutamate receptor activity, monoatomic ion channel activity and metal ion transmembrane transporter activity. Each enriched subclustered pathway is depicted by the respective colours and corresponding genes’ adjusted p-value.

The top five up- and downregulated pathways in microglia, astrocytes and oligodendrocytes showed dysregulation in glial functions, axonogenesis and neuronal support, lipid metabolism, histone/chromatin, mRNA processing and translation/protein folding pathways (**Fig. 5c**). We observed a variable direction of change across all three cell types in glial cell activation, differentiation, axonogenesis and myelin sheath. Histone modification, chromatin remodelling, transcription coregulator activity, and protein folding were exclusively upregulated in all glial cells. Microglia in particular showed significant upregulation in pathways relating to inflammation, histone and chromatin, and mRNA processing. The top five upregulated ORA GO pathways in microglia were subclustered as a network connecting enriched pathway terms with genes annotated with each specific pathway term: cellular response to biotic stimulus, mRNA metabolic process, histone modification, chromatin remodelling, and transcription coregulator activity (**Fig. 5d**).

### 3.5 Bulk proteomics in brain reveals downregulation in translation and upregulation in synaptic pathways

To complement our scRNA sequencing findings in this model, we performed untargeted bulk proteomics in the same setting. Principal component analysis (PCA) of differential protein abundance revealed distinct separation between HF+/SIS+/PIC+ and HF-/SIS-/PIC- controls (n = 3/group) (**Suppl. Fig. S7a**). Of 7859 statistically significant (FDR <0.05) proteins, 2842 were downregulated and 3216 were upregulated in HF+/SIS+/PIC+ offspring relative to HF-/SIS-/PIC- controls (**Suppl. Fig. S7b**). We performed ORA GO analysis based on these differentially expressed proteins (DEPs). Top 10 upregulated pathways included synaptic function and plasticity terms such as presynaptic membrane, CA1 synapse, postsynaptic density and synaptic vesicle (**Fig. 5e**). While top 10 downregulated pathways were predominantly related to ribosome biogenesis and mRNA translation (**Fig. 5f**).

We explored the most enriched upregulated ‘presynaptic membrane’ pathway further by clustering the genes in this pathway into themes based on GO molecular function (**Fig. 5g**). There were five themes that emerged in the genes enriching this pathway including SNARE binding (*Snap25*, *Stx1a*, *Unc13a*, *Snph*), Signalling receptor binding (*Nrxn1*, *Nrxn2*, *Slc6a3*), Monoatomic ion channel activity (*Cacna1c*, *Kcna1*, *Kcna2*, *Kcna3*, *Psen1*), Glutamate receptor activity (*Grin1*, *Gria2*, *Gria3*) and Metal ion transmembrane transporter activity (*Slc6a1*, *Atp1a3*, *Atp2b1*).

### 3.6 scRNA sequencing in peripheral blood leukocytes reveals dysregulation in immune, mRNA processing and translation pathways

UMAP analysis of these samples revealed 12 distinct cell type clusters (**Fig. 6a**). Statistically significant (FDR <0.05) DEGs ranged from 13 to 873 across cell types in HF+/SIS+/PIC+ versus HF-/SIS-/PIC- offspring. Here, we observed the largest proportion of DEGs in neutrophils, macrophages and naïve B Cells (**Fig. 6b**). Next, we performed ORA GO enrichment analysis and we restricted our focus to these 3 cell types based on these DEGs.

**Figure 6.**
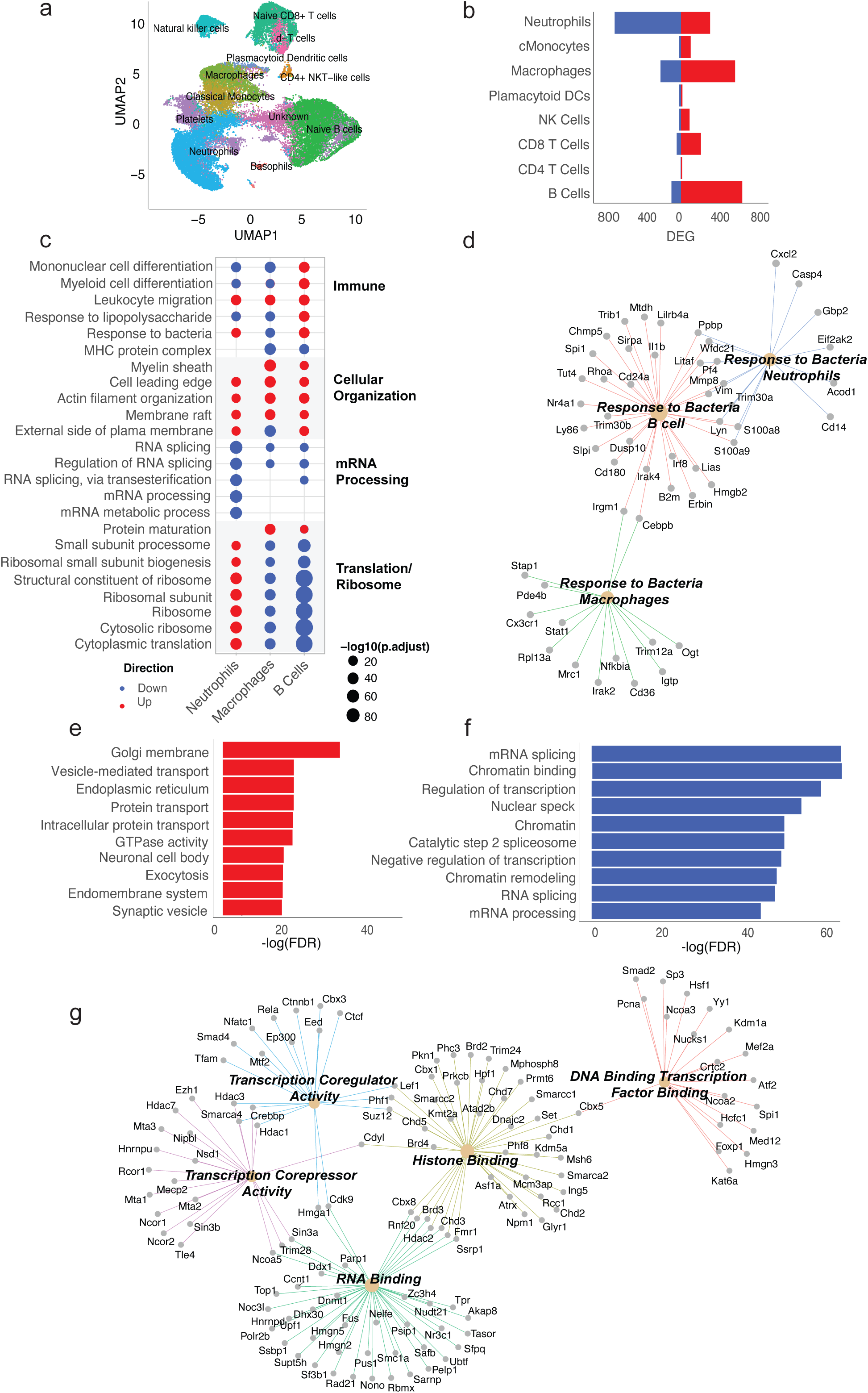
Peripheral blood transcriptome and proteome in male HF+/SIS+/PIC+ offspring versus controls. (**a**) UMAP (uniform manifold approximation of projection) of single-cell populations captured for sequencing: basophils, CD4+ NKT-like cells, classical monocytes, macrophages, naïve B cells, naïve CD8+ T cells, natural killer cells, neutrophils, plasmacytoid dendritic cells, platelets, unknown, ɣ-δ-T cells (**b**) Proportions of significantly differentially expressed genes (DEGs) (FDR <0.05) across leukocyte cell types. Neutrophils (873) display largely downregulated DEGs, macrophages (682) and naïve B cells (647) display largely upregulated DEGs. All three cell types display largest proportions of DEGs. Colours indicate DEG direction of change (*red*: up; *blue*: down). (**c**) Dot plot displaying GO analysis of DEGs (FDR <0.05) HF+/SIS+/PIC+ versus HF-/SIS-/PIC- controls in select cell types (neutrophils, macrophages, B cells) presented on x-axis. Enriched pathway terms are displayed on left y-axis and clustered biological themes of like terms presented on right y-axis. Size of each coloured dot is indicative of the negative log10 of enriched pathways adjusted p-value. The colour of each dot is indicative of the direction of change. (**d**) CNET plot of linked genes from enriched ‘response to bacteria’ GO pathway in the transcriptome in macrophages (downregulated), naïve B cells (upregulated) and neutrophils (upregulated). (**e**) Bar plot of bulk proteome depicting top 10 enriched GO pathways amongst proteins with increased abundance. Statistical significance depicted by negative log(FDR). (**f**) Bar plot of bulk proteome depicting top 10 enriched GO pathways amongst proteins with decreased abundance. Statistical significance depicted by negative log(FDR). (**g**) Subclustered CNET plot of the downregulated GO pathway ‘chromatin binding’ in proteome. Subclusters are based on molecular function and include: transcription coregulator activity, transcription corepressor activity, RNA binding, histone binding and DNA binding transcription factor binding. Each enriched subclustered pathway is depicted by the respective colours and corresponding genes’ adjusted p-value.

Top five up- and downregulated pathways in neutrophils, macrophages and naïve B cells showed dysregulation in immune, cellular organization, mRNA processing, and ribosomal/translational pathways (**Fig. 6c).** We observed upregulated cellular organization including myelin sheath, actin filament organization, and downregulated mRNA processes including RNA splicing, mRNA metabolic process, across all three cell types. However, immune pathways including mononuclear cell differentiation, response to lipopolysaccharide, response to bacteria, and translational pathways showed variable directions. Neutrophils exhibited mixed regulation in immune pathways while showing upregulation in ribosomal and translational pathways. In contrast, naïve B cells predominantly revealed upregulated immune pathways but downregulated ribosomal and translational pathways.

The response to bacteria pathway was upregulated in neutrophils and naïve B cells, but downregulated in macrophages, with different genes enriching this pathway (**Fig. 6d**). Genes which enriched the response to bacteria pathway included the following in neutrophils (*Cd14, Casp4, S100a8, S100a9)*, macrophages (*Irak2, Nfkbia, Cx3cr1*) and naïve B cells (*Il1b, S100a8, S100a9, Irak4, Irf8)*.

### 3.7 Peripheral blood proteomics reveals downregulated chromatin and mRNA processing pathways

We also performed bulk proteomics in peripheral blood leukocytes in the same setting. PCA analysis of differential protein abundance revealed distinct separation between HF+/SIS+/PIC+ and HF-/SIS-/PIC- controls (**Suppl. Fig. S7c**). Of 4464 statistically significant (FDR <0.05) proteins, 2720 were downregulated and 3134 were upregulated in stressed offspring relative to controls (**Suppl. Fig. S7d**). We performed ORA GO analysis based on these DEPs. Top 10 upregulated pathways included intra- and extracellular transport functions such as Golgi membrane, vesicle-mediated transport, synaptic vesicle, protein transport, and GTPase activity (**Fig. 6e**). Top 10 downregulated pathways were predominantly related to splicing and epigenetic modifications such as, chromatin binding/remodelling, and mRNA processing **(Fig. 6f).**

We explored this highly downregulated chromatin binding pathway further by clustering the genes in this pathway into themes based on GO molecular function (**Fig. 6g).** There were five themes that emerged and the genes enriching these pathways included histone binding (*Hdac2, Kmt2a, Kdm5a, Chd1, Chd2, Chd3, Chd7, Brd2, Brd3, Brd4*), transcription coregulator activity (*Ep300, Rela*), transcription corepressor activity (*Hdac1, Hdac3, Hdac7*), DNA binding transcription factor binding (*Kdm1a, Kat6a*), and RNA binding (*Fmr1, Dnmt1, Hmgn2, Hmgn5*).

## 4. Discussion

In this study, we explored the behavioural and molecular impacts of cumulative environmental stress exposures on foetal neurodevelopment into adulthood. Furthermore, we aimed to identify transcriptomic and proteomic alterations in the brain and periphery to uncover overlapping interactions in the pathogenesis of NDDs.

We demonstrated that cumulative environmental exposures impact social behaviours in male HF+/SIS+/PIC+ offspring, a phenotype which aligns with one of the core symptoms of autism and multifactorial aetiology of NDDs. In addition, triple-hit males displayed the highest NDD-behavioural risk profile compared to non-stressed controls. While we did not perform analyses between single-hit versus triple-hit groups due to small sample sizes and statistical limitations, we hypothesise that larger sample numbers may capture significant effects in single-hit or double-hit models. Nonetheless, we propose that these results are translationally meaningful, as epidemiological evidence demonstrate that cumulative maternal and postnatal environmental influences markedly increase the risk of autism and ADHD (Becker, 2007; Bilbo et al., 2018; Han et al., 2021b; Nielsen et al., 2024; Volk et al., 2014). In contrast, we did not observe any social deficits in males exposed to only mSIS or postnatal poly(I:C). Interestingly, HF+/SIS-/PIC- offspring displayed normal social recognition but displayed sociability deficits comparable to our triple-hit group. These results align with a recent study showing mHFD-induced MIA impaired sociability in male offspring via dysregulated tryptophan metabolism (Sun et al., 2025). Here, we presume that chronic mHFD exposure poses as a significant developmental modulator in the face of additional environmental exposures in early life. Furthermore, these results emphasise the clinical significance of poor maternal diet during pregnancy and the neurodevelopmental consequences of chronic exposure.

While not the main focus of this study, there appeared to be a sex-specific vulnerability to these behavioural deficits, as we did not detect any alterations to social behaviours in female HF+/SIS+/PIC+ offspring (**Suppl. Fig. S4**) compared to sex-matched controls or an abnormal NDD-behavioural risk profile compared to male HF+/SIS+/PIC+ offspring. The sexual dimorphism underlying NDDs is highly nuanced; in the context of autism, there are two contrasting theories concerning the underdiagnosis of females. The ‘female protective effect’ suggests that females require a greater etiological load to exhibit the same behavioural symptoms as males, while the ‘female autism phenotype’ posits that female symptom presentation does not align well with current diagnostic criteria, leading to masking or ‘camouflaging’ (Hull et al., 2020; Robinson et al., 2013). While we could not address these complexities here, they warrant further exploration in a context of cumulative environmental exposures.

Given the complexity of NDD pathophysiology, we analysed the transcriptome and proteome of HF+/SIS+/PIC+ male offspring to identify key biological processes disrupted by multiple environmental exposures at a systems level. Furthermore, as peripheral blood is the most accessible tissue in clinical settings, we characterised blood -omics profiles alongside brain tissue to enhance the model’s translational relevance and identify whether these molecular alterations are conserved between both compartments. In the transcriptome of microglia, astrocytes and oligodendrocytes, we observed a consistent theme of immune-related dysregulation, particularly in microglia. Of the top five upregulated pathways in these cells, genes that enriched for the *cellular response to biotic stimulus* pathway are all implicated in regulating inflammation, neurodevelopment and microglial activity (e.g., *Il1a, Irak4, Notch1*) (Lombardo et al., 2018; Lv et al., 2017; Sreenivas et al., 2024; Zhang et al., 2023). Our pathway enrichment analysis in the transcriptomes of neutrophils, macrophages and naïve B cells revealed immune pathways to be dysregulated, with both up- and downregulation. Moreover, the direction of change varied between innate and adaptive immune cells. Peripheral macrophages displayed a distinct set of genes that enriched for the downregulated expression of the response to bacteria pathway in HF+/SIS+/PIC+ male offspring. The CX3C chemokine receptor 1 (*Cx3cr1*) gene particularly stands out, as the significance of this G protein-coupled receptor is primarily restricted to microglial functions. Appropriate CX3CR1 activation is required for microglia-mediated synaptic pruning during early development (Estes and McAllister, 2015) and animal models deficient in this receptor display autism-like behaviours (Zhan et al., 2014). Its dysregulated expression in human peripheral blood and post mortem brain tissue has also been shown to be highly associated with NDDs such as schizophrenia (Bergon et al., 2015) and ASD (Ishizuka et al., 2017; Suzuki et al., 2013b; Voineagu et al., 2011). Thus, its dysregulation in peripheral macrophages here highlights neuro-immune interactions that are implicated in the pathogenesis of NDDs. Taken together, these transcriptomic alterations point to systemic immune dysregulation as one of many consequences of cumulative environmental exposures in early life. Moreover, these findings complement existing evidence suggesting chronic immune dysregulation between the periphery and the brain may be associated with NDDs (Ashwood et al., 2004; Enstrom et al., 2010; Estes and McAllister, 2015; Martino et al., 2020) and highlight peripheral immune and glial priming as key mechanisms warranting further investigation.

Beyond immune dysregulation, ribosomal biogenesis, translation and mRNA processing were additional functional pathways we found to be disturbed across the periphery and CNS. The transcriptome of peripheral blood macrophages and naïve B cells revealed *cytoplasmic translation* and *ribosome* associated functions to be significantly downregulated, whereas neutrophils showed upregulation. In all brain glia cells, pathways such as *mRNA processing* and *translation* were exclusively upregulated. Dysregulated translational control and disrupted proteostasis have emerged as further disease promoting mechanisms associated with NDD risk. The regulation of proteostasis is important for maintaining synaptic homeostasis, as balanced synaptic protein turnover is crucial for the remodelling of synapses (Alvarez-Castelao and Schuman, 2015; Chen et al., 2019; Kalish et al., 2021).

As synaptic impairments are a common pathological hallmark of NDDs (Ben-David and Shifman, 2012; Bourgeron, 2015; Brandt et al., 2014; Ugarte et al., 2023), we were intrigued to find evidence of synaptic protein alterations in the proteome of male HF+/SIS+/PIC+ offspring. In the brain, proteins with significantly increased abundance showed enriched GO pathways involved in synaptic structure and function. Further, we observed *translation at presynapse* to be a pathway enriched in proteins with significantly reduced abundance. This was consistent with our findings of dysregulated translation and ribosome functions in the transcriptome and thus, emphasises the significance of appropriate regulation of synaptic protein turnover in the context of normal neurodevelopment. Moreover, further analysis of the *presynaptic membrane* pathway in proteins with increased abundance in the brain revealed several genes linked to autism and other NDDs, including: *Grin1, Nrxn1, Gria1-3*, and *Cacna1c*. These genes have functional relevance to synaptogenesis, glutamate receptor activity, ion channel activity or receptor binding – processes often dysregulated in NDDs (Bourgeron, 2015; McCarthy and Wright, 2017; Tang et al., 2013; Voineagu et al., 2011). While we did anticipate finding synaptic alterations in the brain proteome, we did not expect this theme to be common to peripheral immune cells as well. Specifically, pathways such as *neuronal cell body, endomembrane system* and *synaptic vesicle* were enriched in proteins of increased abundance in the blood. Collectively, these findings capture impaired synaptic stability and regulation that is evident in the periphery.

Lastly, we observed alterations to epigenetic modifiers and changes to transcriptional regulation in our omics analyses. The transcriptome of brain glia displayed pronounced histone modification and chromatin remodelling pathway upregulation; while that of the peripheral blood displayed downregulation in pathways associated with post-transcriptional regulation. However, proteomic analyses of peripheral immune cells revealed reduced protein expression in pathways strongly associated with chromatin. Interestingly, we observed both the microglial transcriptome, and the peripheral blood proteome shared gene families enriched in histone-associated functions, particularly those within the chromatin remodelling protein (e.g. CHDs) or histone lysine demethylase enzyme (e.g. KDMs) families. Both have influence on various aspects of neuronal development, such as neural progenitor generation, cell-specific differentiation and expansion, migration and circuit integration (Hsieh and Gage, 2005). In a small, but important percentage of cases, mutations in chromatin remodelling genes and histone demethylases appear to be overrepresented in children with NDDs (Borroto et al., 2024; De Rubeis et al., 2014; Mossink et al., 2021). In this model, we observed common post-transcriptional mechanisms to be evident following multiple environmental stress exposures in the absence of genetic predisposition. However, further studies are needed to dissect how epigenetic modifiers differentially impact brain and blood homeostasis following early life adversity.

We acknowledge several limitations in this study. First, we recognise that our behavioural analyses were limited in statistical power due to low sample size in our mHFD exposed offspring. While translationally relevant, exposing dams to multiple stressors and maintaining them on a high-fat diet for prolonged periods significantly affects reproductive outcomes (Bucknor et al., 2024). For future longitudinal studies modelling multiple environmental stressors in mice, it is important to note a large number of females and resources are necessary to generate sufficient numbers of offspring of both sexes. This will also allow for controlling for litter effects, which we were unable to address in this study. In addition, while we did not observe any profound behavioural impairments in females at 12 weeks of age, we acknowledge that we cannot fully conclude that females are protected from cumulative stress exposures. Rather, we hypothesise that there could either be a delay in the onset of detectable behavioural changes in females or they may present differently. Therefore, extending upon our existing behavioural battery to include an additional time of testing and further NDD-associated tests [for review, see: (Silverman et al., 2010)], would be valuable for understanding how these stressors influence female behaviour. Additionally, previous reports have demonstrated that females do exhibit unique transcriptional reprogramming in the brain following early developmental adversity (Glendining et al., 2018; Kalish et al., 2021), highlighting the need for further -omics profiling in female offspring subjected to multiple exposures that we were unable to illustrate in the current study. Lastly, this study was not designed to establish biological causality between cumulative stress-induced behavioural changes and NDD-associated molecular mechanisms. Future studies exploring causal mechanisms underlying multiple environmental exposures and NDD pathogenesis will be crucial for prevention and the development of biomarker-specific therapies.

Overall, we conclude that the cumulative effects of multiple early-life environmental exposures persist into adulthood and in a sex-specific manner. In males, we identified overlapping peripheral-CNS immune, ribosomal and epigenetic alterations that are associated with autism-like social impairments. We speculate that these behavioural changes may be a consequence of maternal stress-induced epigenetic priming in peripheral immune cells and glia *in utero*, which significantly increase disease susceptibility following postnatal poly(I:C) exposure. The convergent molecular impairments we observed between the periphery and the brain should be characterised further in a cumulative context and open the potential of peripheral blood to serve as a reliable prognostic and diagnostic biomarker for NDDs.

## Data availability

Raw FASTQ files associated with the scRNA sequencing data from peripheral blood and brain cells described here are accessible through NCBI’s Gene Expression Omnibus (GEO) (accession GSE289585).

Reviewer access details:

To review GEO accession GSE289585:

Go to https://www.ncbi.nlm.nih.gov/geo/query/acc.cgi?acc=GSE289585 Enter token spediwewrlcrvyh into the box

The mass spectrometry proteomics data for brain samples have been deposited to the ProteomeXchange Consortium via the PRIDE partner repository with the dataset accession: PXD060752 and DOI:10.6019/PXD060752.

Reviewer access details:

Log in to the PRIDE website using the following details:

Project accession: PXD060752

Token: t1FymYi8xMjY

Alternatively, reviewer can access the dataset by logging in to the PRIDE website using the following account details:

Username

Password: OfLET3KNpQYj

The mass spectrometry proteomics data for blood samples have been deposited to the ProteomeXchange Consortium via the PRIDE partner repository with the dataset accession PXD060609 and DOI:10.6019/PXD060609.

Reviewer access details:

Log in to the PRIDE website using the following details:

Project accession: PXD060609

Token: kUHHPEEJP2pM

Alternatively, reviewer can access the dataset by logging in to the PRIDE website using the following account details:

Username

Password: GPO0wB0kNrQb

## Acknowledgements

The authors acknowledge the Animal Behavioural facility and the assistance of Laboratory Animal Services (LAS) at The University of Sydney. We also thank representatives from Specialty Feeds (WA, Australia) for their expert advice on all diet related queries during the study. We also thank the team at Australian Genomics Research Facility (AGRF) for scRNA-sequencing and Children’s Medical Research Institute (CMRI) for bulk proteomics.

## Declaration of generative AI in scientific writing

During the preparation of this work the author(s) used ChatGPT in order to improve readability and concise language. After using this tool/service, the author(s) reviewed and edited the content as needed and take(s) full responsibility for the content of the publication.

**Appendix A. Supplementary data**

**Appendix B. Extended methods**

