## Appendix A - Supplementary Data for "Cumulative pregnancy and postnatal environmental exposures impact social behaviour in male mice associated with epigenetic, ribosomal, and immune dysregulation"

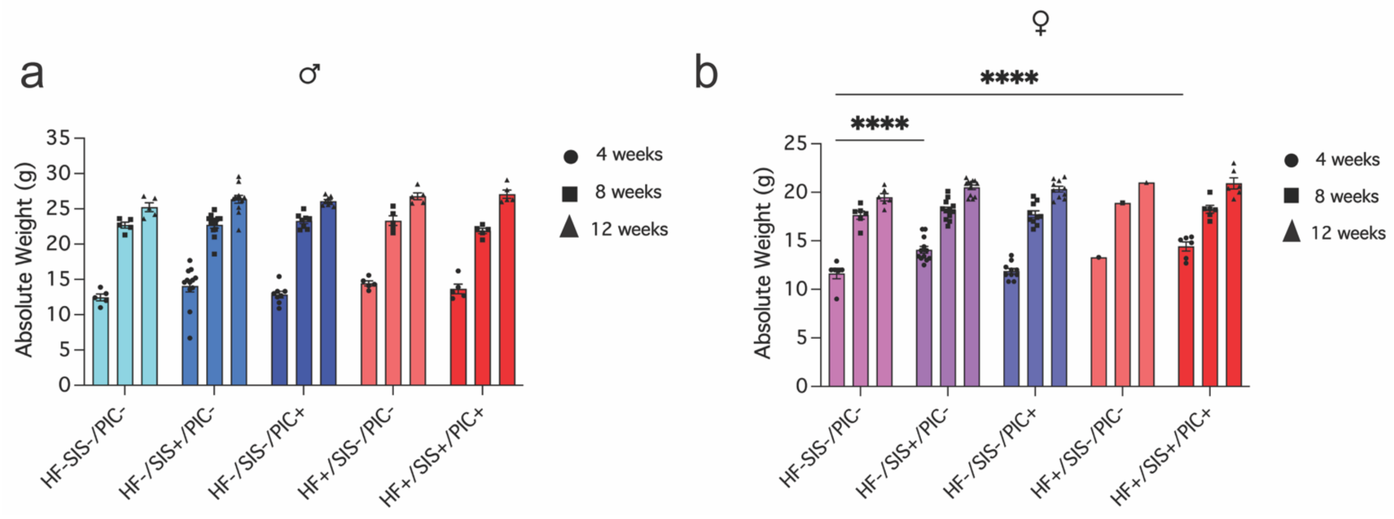


**Figure S1. Offspring demonstrate normal physiological growth into adulthood across stress groups.** (**a**) *males:* Absolute weight (g) across all groups at 4, 8 and 12-weeks of age. Mixed effects model followed by type III ANOVA. Main effect of group (ns); age (****) $F_{2,60} = 1368$, $p =< 0.0001$; group x age (*) $F_{8,60} = 2.43$, $p = 0.023$. Followed by Dunnett's post-hoc test (ns). (**b**) *females:* Absolute weight (g) across all groups at ages described above. Mixed effects model followed by type III ANOVA. Main effect of group (**) $F_{4,30} = 5.2$, $p = 0.002$; age (****) $F_{2,60} = 305.2$; group x age (**) $F_{8,60} = 3.34$, $p = 0.003$. Followed by Dunnett's post-hoc test:4 weeks of age (HF-/SIS-/PIC- vs HF-/SIS+/PIC- and HF+/SIS+/PIC+ (****, $p<0.0001$)). Data shown as mean ± SEM; mixed effects model. Fixed effects: group, age; random effect: mouse. Legend -males (♂) and females (♀).

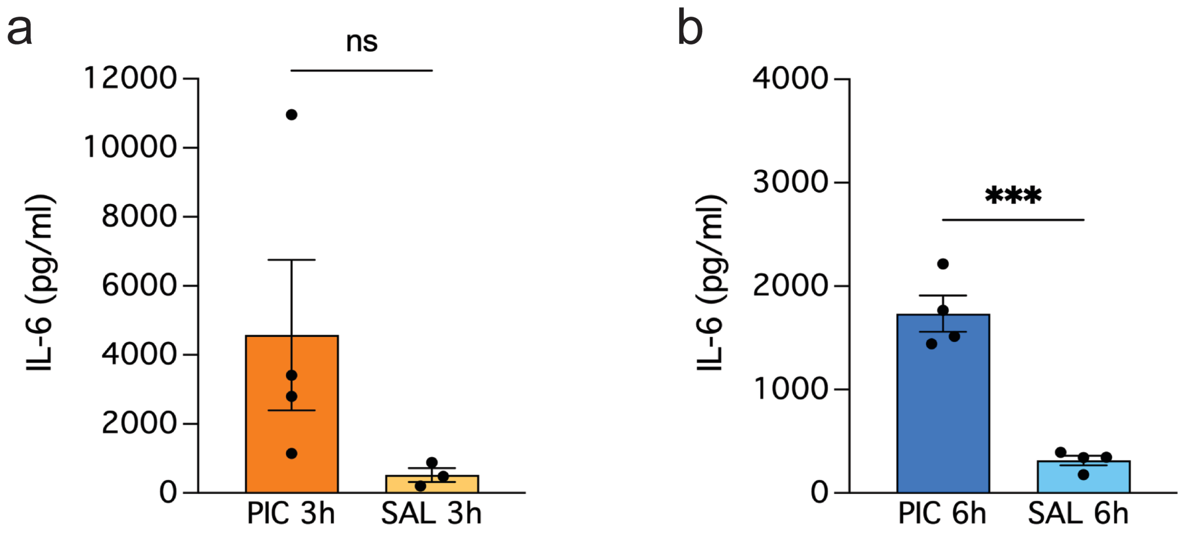


**Figure S2. Poly(I:C) stimulation increases peripheral plasma levels of IL-6 at 3 and 6hrs.** (**a**) Plasma IL-6 (pg/ml) concentration 3 hours post- poly(I:C) intraperitoneal injection and/or saline vehicle. Mann-Whitney test (p = 0.0571); *ns*. (**b**) Plasma IL-6 (pg/ml) concentration 6 hours post- poly(I:C) intraperitoneal injection and/or saline vehicle. Unpaired t-test (***p = 0.0002). Data shown as mean ± SEM. PIC = poly(I:C), SAL = saline, ns = not significant.

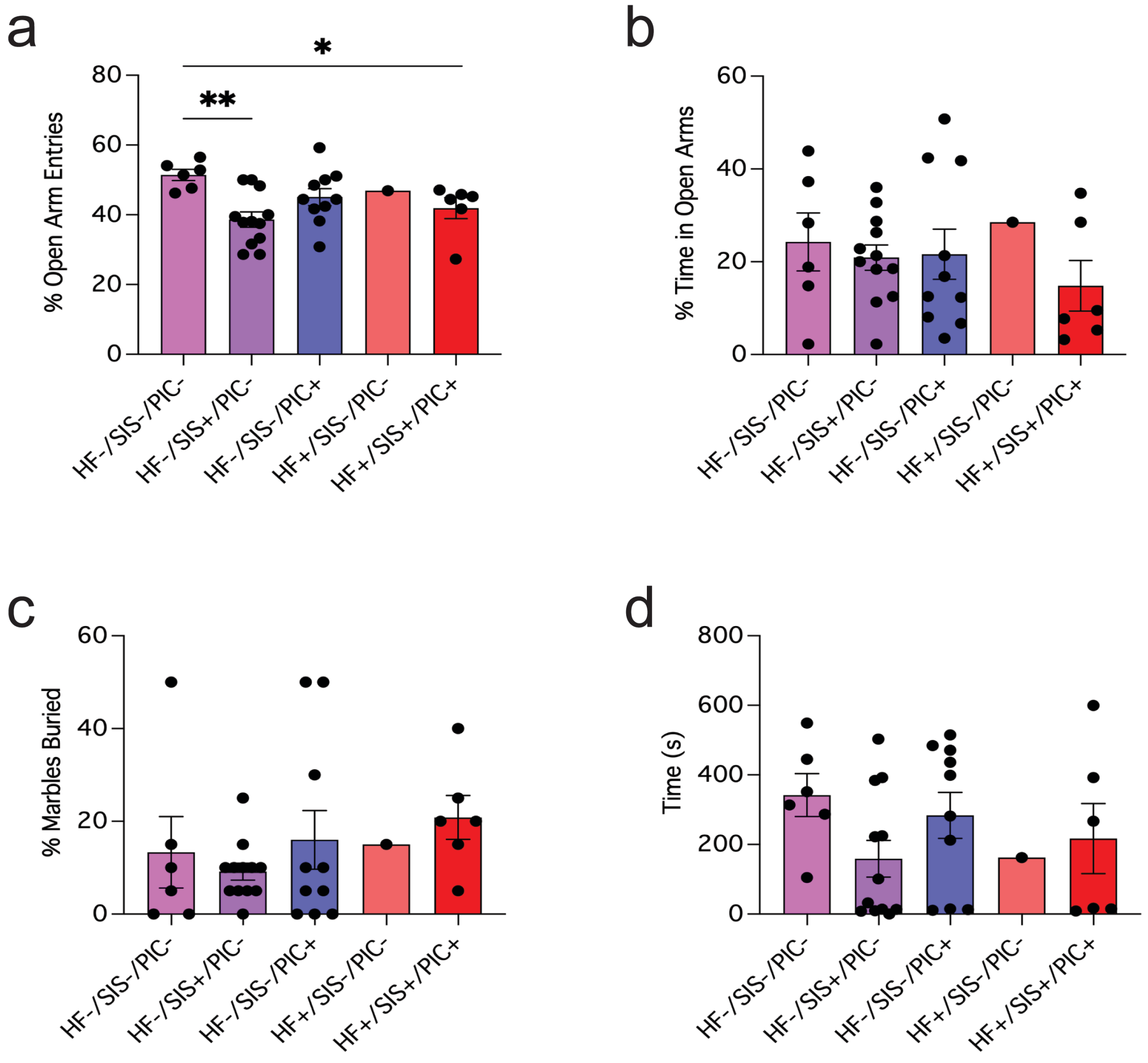


**Figure S3. Female offspring exposed to mSIS alone and all three stress factors display a reduction in open arm entries of the plus maze.** (**a-b**) Elevated plus maze, (**a**) percentage of open arm entries. Ordinary one-way ANOVA. Main effect of group (*) F4,36 = 3.64, p = 0.0137. Followed by Dunnett's post-hoc test: (HF-/SIS-/PIC- vs HF-/SIS+/PIC- and HF+/SIS+/PIC+ (**, p = 0.0031) and (*, p = 0.035), respectively. (**b**) percentage of time spent in the open arms; (**c**) marble burying test; (**d**) time spent grooming. Data shown as mean ± SEM.

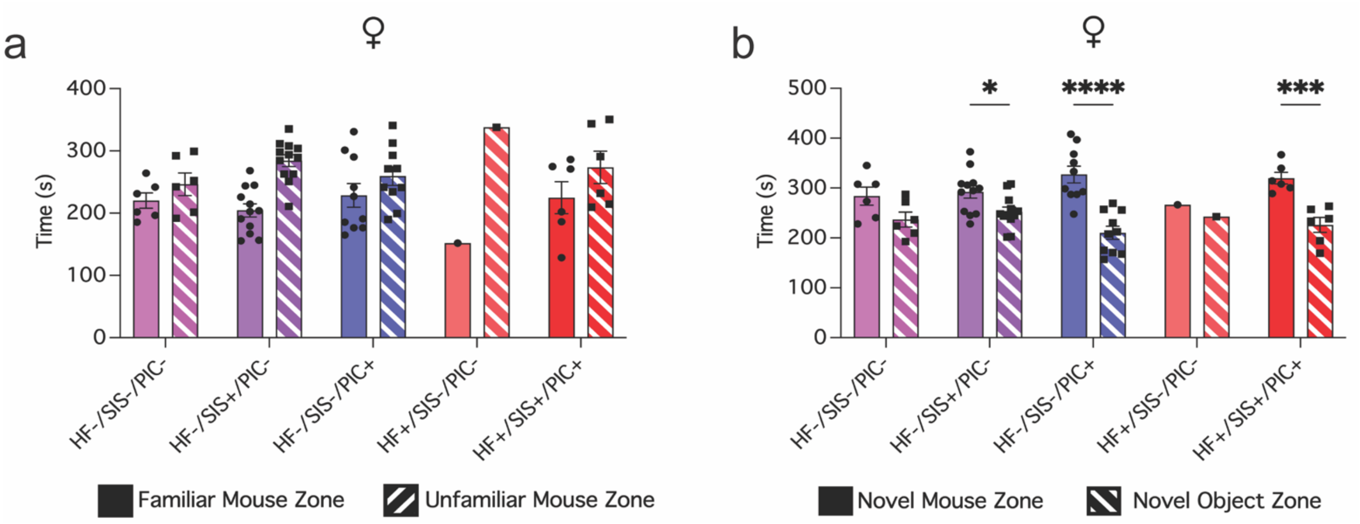


**Figure S4. Females across all stress groups do not display social impairments.** (**a-b**) 3-chamber social preference test, (**a**) time (s) spent in familiar mouse zone and unfamiliar mouse zone (**b**) time (s) spent in novel mouse zone and novel object zone. Linear model followed by Type III ANOVA. Main effect of group (ns), zone (ns), group x zone (*) F4,60 = 3.07, p = 0.022. Followed by Tukey's post-hoc test for within group comparisons between novel mouse zone versus novel object zone: (HF-/SIS-/PIC+ (****,p<0.0001), HF-/SIS+/PIC- (*, p = 0.0201) and HF+/SIS+/PIC+ (***, p = 0.0002). Data shown as mean ± SEM.

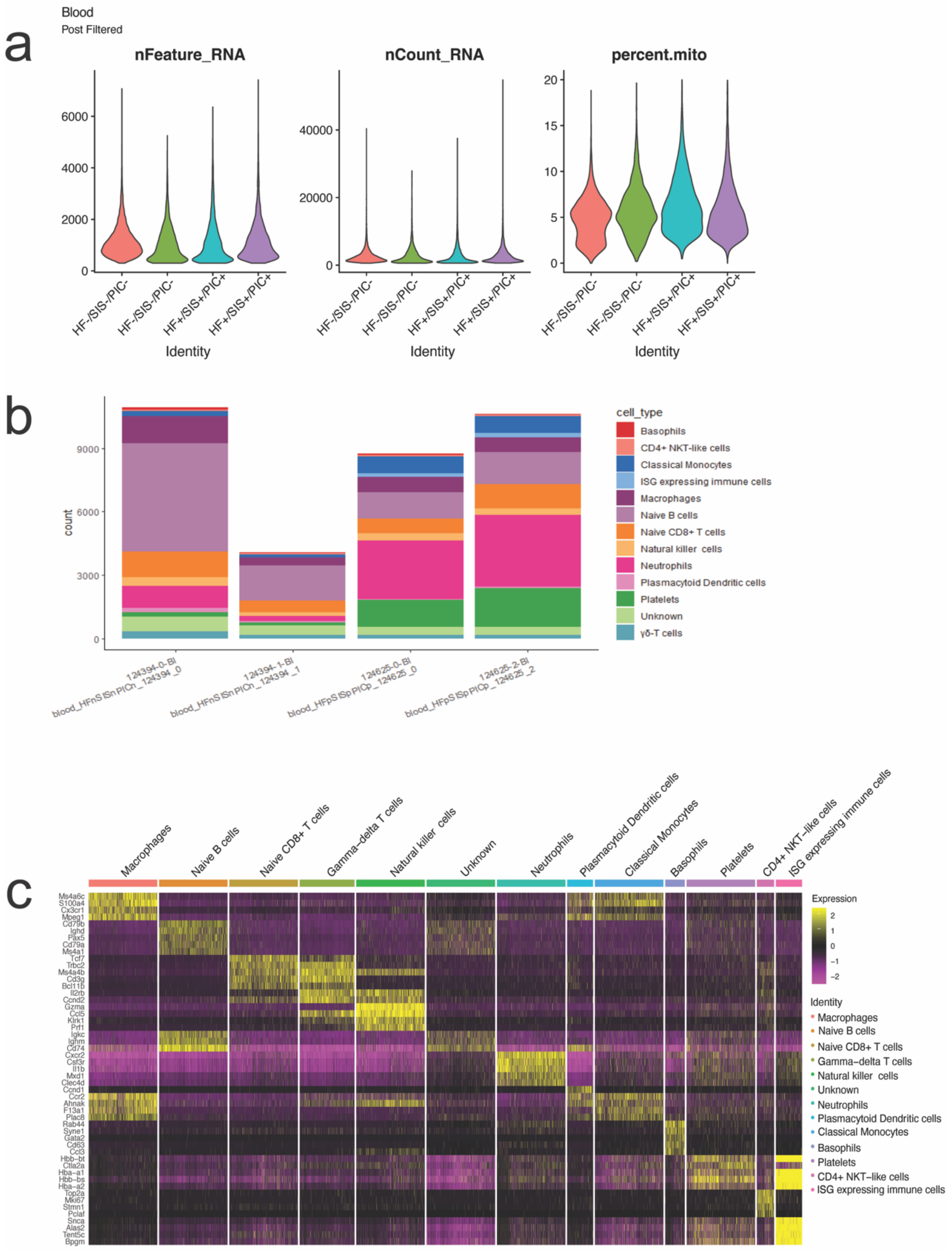


**Figure S5. scRNA QC summary and heatmap of top differentially expressed genes by cell type for peripheral blood leukocytes.** (**a**) Peripheral blood isolated from male HF+/SIS+/PIC+ and HF-/SIS-/PIC- controls. nFeature_RNA is indicative of the number of genes detected per cell, nCount_RNA is the number of transcripts per cell and percent.mito is indicative of the amount of mitochondrial contamination from dead or dying cells. (**b**) Cell proportions (counts) across samples (not scaled). (**c**) Heatmap displaying top differentially expressed genes per cell type. Data from n = 2 mice / group.

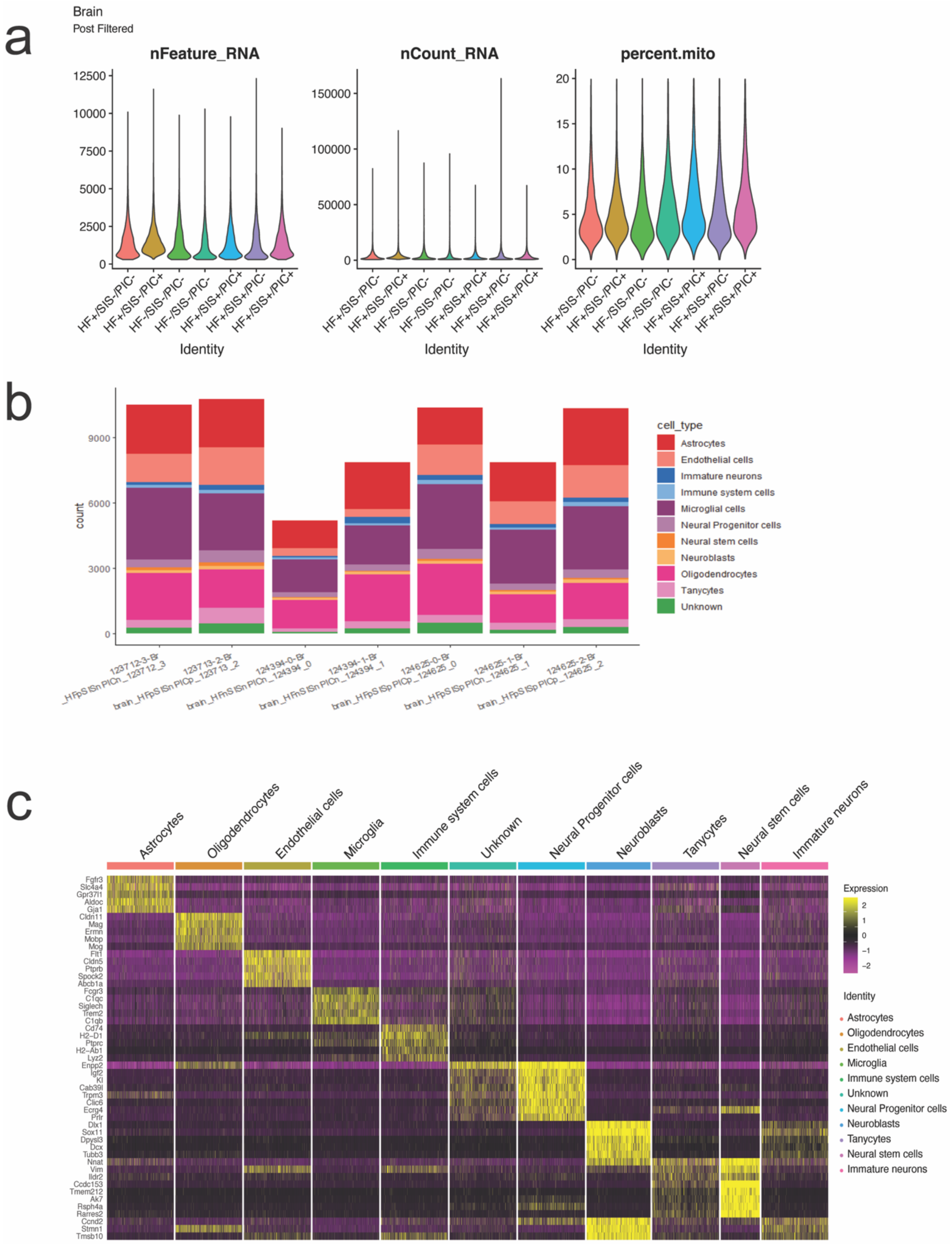


**Figure S6. scRNA QC summary and heatmap of top differentially expressed genes by cell type for brain cells.** (**a**) Brain cells isolated from HF-/SIS-/PIC-, HF+/SIS+/PIC+ and intermediate male samples that were used for pilot data not described in this manuscript (HF+/SIS-/PIC-, HF+/SIS-/PIC+ and HF+/SIS+/PIC-). nFeature_RNA, nCount_RNA and percent.mito are displayed for number of genes detected per cell, transcripts per cell and proportion of mitochondrial contamination from dead or dying cells. (**b**) Cell proportions across samples (not scaled). (**c**) Heatmap displaying top differentially expressed genes per cell type. Data from n = 2 mice / group between HF+/SIS+/PIC+ and HF-/SIS-/PIC- male controls. Intermediate samples n = 1 / group.

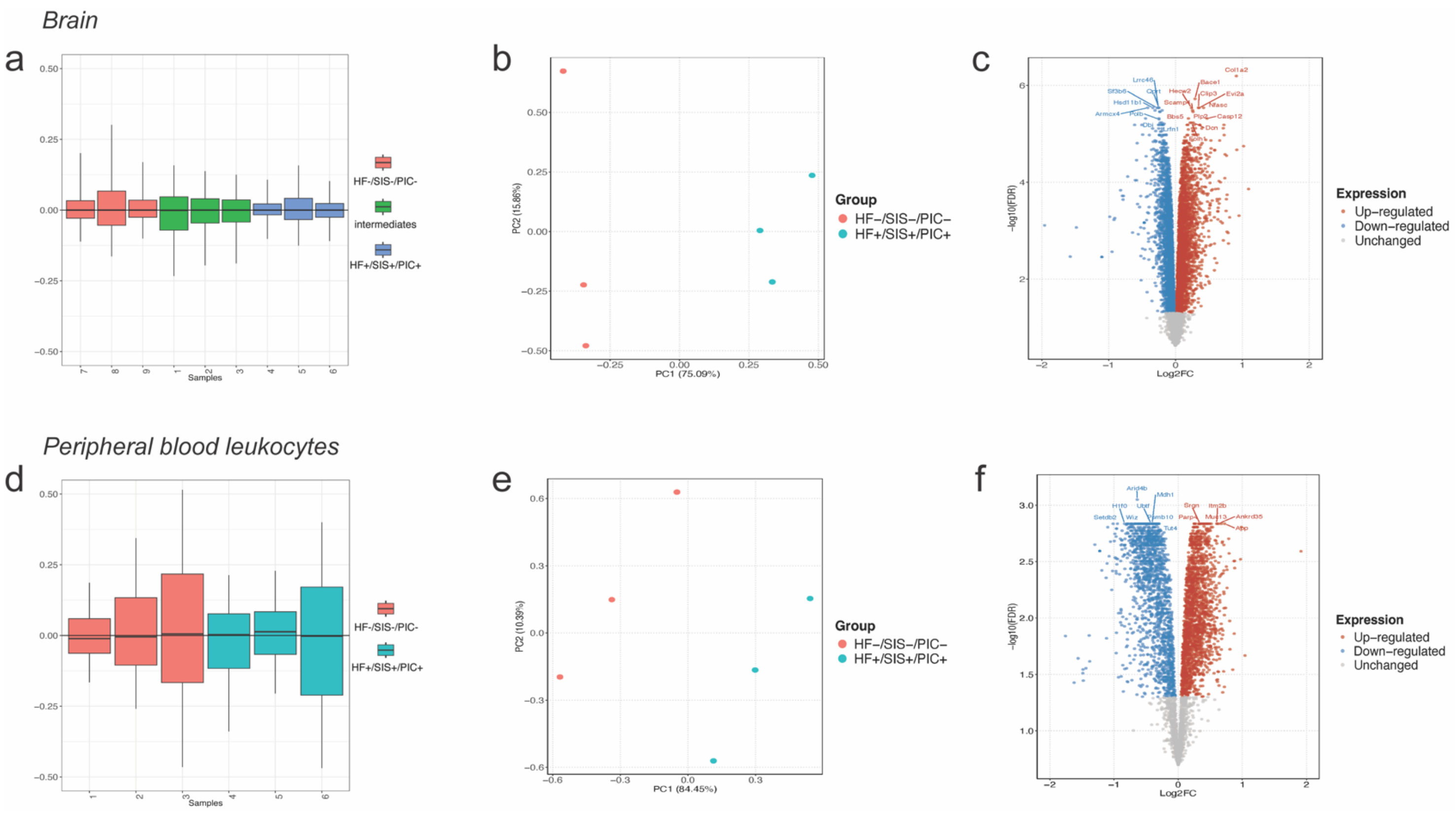


**Figure S7. Proteomics RUV normalisation, principal component analysis (PCA) plots and differentially expressed proteins (DEPs).** (**a**) Box and whiskers relative log expression plot of brain samples after RUV normalisation. Intermediates samples were not described in this manuscript. (**b**) PCA of proteomic data depicting two distinct clusters of stressed (HF+/SIS+/PIC+) versus control (HF-/SIS-/PIC-) offspring in brain tissue. (**c**) Volcano plot depicting top differentially abundant proteins. Coloured dots indicate statistical significance (FDR<0.05). Positive $\log_2$ fold change (Log2FC) (red) illustrates up-regulated expression (e.g. Col1a2, Bace1, Clip3, Evi2a, Nfasc, Casp12, Plp2, Dcn, Folh1, Bbs5, Scamp4, Hecw2), and negative $\log_2$ fold change (blue) illustrates down-regulated expression (e.g. Lrrc46, Qprt, Lrfn1, Dbi, Polb, Hsd11b1, Sf3b6) in stressed males relative to control males. (**d**) Box and whiskers relative log expression plot of blood samples after RUV normalisation. (**e**) PCA of proteomic data depicting two distinct clusters of stressed (HF+/SIS+/PIC+) versus control (HF-/SIS-/PIC-) offspring in peripheral blood leukocytes. (**f**) Volcano plot depicting top differentially abundant proteins. Coloured dots indicate statistical significance (FDR<0.05). Positive $\log_2$ fold change (Log2FC) (red) illustrates up-regulated protein expression (e.g. Srgn, Itm2b, Muc13, Parp4, App, Ankrd35) and negative $\log_2$ fold change (blue) illustrates down-regulated protein expression (e.g. Arid4b, Mdh1, Psmb10, Tut4, Wiz, H1f0, Setdb2, Ubtf) in stressed males relative to control males. Data from n = 3 mice / group.
