## Appendix B - Extended Methods for "Cumulative pregnancy and postnatal environmental exposures impact social behaviour in male mice associated with epigenetic, ribosomal, and immune dysregulation"

#### Maternal stress model study design and dietary information

Details and experimental findings from our maternal stress model have been previously described here (Bucknor et al., 2024). Briefly, maternal stress groups involved chronic exposure to either a high-fat diet (HF+) or a control low-fat diet (HF-) lasting before and during pregnancy, including the postpartum lactation period. In addition, from each diet group, mice were randomly assigned to either control conditions or psychosocial stress exposure, which entailed 6-weeks of social instability stress prior to inducing pregnancy. Therefore, 4 maternal stress groups were generated:

1) HF+/SIS+
2) HF+/SIS-
3) HF-/SIS+
4) HF-/SIS-

#### On-site breeding of offspring

Breeding of offspring was performed in house in a specific pathogen free environment (SPF) at the Charles Perkins animal facility (20-24℃, 40-70% humidity, 12h light:dark cycle) (University of Sydney, Sydney Australia). Female oestrus cycle was stimulated 3 days prior to their first mating encounter to increase receptivity to mating. This was done by placing male urine and bedding into female cages. Mating design was both 1:2 and 1:1 breeding pairings. Females were housed with a male and monitored for weight gain from gestation day (GD)0 – 13. Females that gained >3g in body weight were considered pregnant and single-housed till birth. Females that did not conceive after their first mating encounter, were re-paired with a new male a week later. Each female (n = 45) was subjected to a maximum of two mating encounters. Out of 45 females bred, 21 females conceived and carried litters to full term used in this study. Males (n = 11) were used for multiple breeding rounds and their exposure to either the high-fat or low-fat diet was consistent between rounds (i.e., a subset of males bred with high-fat diet fed females only or low-fat diet fed females only).

#### Offspring study design and dietary information

Offspring were generated via-on-site breeding from each maternal stress group as described above (n = 71 total). Immediately after birth and during the lactation period, offspring remained with their mothers undisturbed till weaning (postnatal day 28). At weaning, offspring were sexed, weighed and transitioned to standard grain-based laboratory chow (SF00-100, Specialty Feeds, WA Australia) for the remainder of the study and weighed again at 8- and 12-weeks of age. Mice were group housed with 4-5 mice/cage. The following day (postnatal day 29), offspring were randomised to receive either 10 mg/kg poly(I:C) (high molecular weight, catalogue #: tlrl-pic5, Invivogen) in endotoxin-free 0.9% NaCl or vehicle alone via intraperitoneal injection.

**Poly(I:C) validation**

Poly(I:C) immune activation (PIA) was validated using an independent cohort of male C57Bl/6 mice (n = 15) and the same batch of poly(I:C) was used to avoid batch effects. PIA was defined by an acute increase in plasma IL-6 concentration at 3- and 6-hours post-injection (see **Suppl. Fig. S2**). Plasma concentrations of IL-6 was determined using a commercial IL-6 ELISA kit (ab22503; sensitivity = 11.3 pg/mL; Abcam) according to manufacturer's instructions. Plasma samples were diluted 1:8 and analysed in duplicate. Absorbance was measured at 450nm using a Tecan infinite M1000 Pro plate reader. Concentrations of IL-6 were calculated using the AssayFit Pro 96 well curve fitting ELISA calculator (https://www.assayfit.com/home.html). Standard curve was determined by fitting the data to a 4-parameter logistic curve (4PL).

**Behavioural testing**

Behavioural testing began when male and female offspring reached 12-weeks of age and carried over a 2-week period (mice were subject to a maximum of two tests per week). All tests were performed during the light cycle between 0900 and 1700h. The behavioural battery of tests selected have relevance to behavioural symptoms observed in neurodevelopmental disorders (Silverman et al., 2010). Testing was performed in order of most to least stressful, starting with the elevated plus maze and 3-chamber social preference test, and ending with marble burying and self-grooming. Mice were habituated in their home cages in a dimly lit procedure room (40-60 lux) adjacent to the testing room for a minimum of 30-minutes prior to testing and given 1-4 days of rest in between individual tests. Mice that were tested were separated from untested cage mates until all cage mates completed testing to prevent stress-induced pheromonal interference (Bind et al., 2013). All apparatuses were cleaned with 70% ethanol between tests, and the experimenter was absent from the room during testing. The elevated plus maze, 3-chamber social preference test and self-grooming behaviours were recorded using a digital webcam (Logitech Brio 4K Webcam). Elevated plus maze and 3-chamber social preference output was analysed using AnyMaze video tracking software (V7.20), while self-grooming was manually scored using Behavioral Observation Research Interactive Software (Boris). Marble burying output was obtained from images taken of the bedding at the end of the test and manually scored by two independent researchers. Detailed descriptions of each test are provided below.

**Elevated plus maze**

The elevated plus maze test was used to assess differences in natural exploratory drive that can inform changes to innate anxiety-like behaviour in rodents. The maze platform was elevated 43 cm off the floor and consisted of two open arms and two enclosed arms (30 cm L x 5.5 cm W). The closed arms were enclosed with dark plexiglass (16 cm H). Mice typically demonstrate an innate preference for the two enclosed arms but will explore the open arms as well; an increase in time spent or number of entries in the open arms is reflective of anti-anxiety behaviour (Silverman et al., 2010; Walf and Frye, 2007). The apparatus was placed in the middle of the test room with a webcam camera mounted directly above the maze; and the lighting in the centre of the maze was brightly lit (250-400 lux). The test mouse was placed gently inside the centre of the maze facing the open arms and allowed to freely explore for a 5-min test period. Parameters measured included total distance travelled, percentage of open arm entries and percentage of time spent in the open arms of the maze.

**3-chamber social preference**

Mice are social animals that engage in high levels of reciprocal social interaction, thus various assays have been established to assess changes in social behaviours. One such assay is the 3-chamber social preference test that evaluates sociability and social novelty over three sequential 10-minute trials (habituation, sociability, social novelty).

In this study, sociability and social novelty were assessed using a rectangular, 3-chambered apparatus made from white polycarbonate under dim lighting (40-60 lux). Each chamber was 20cm (L) x 40cm (W) x 22cm (H). Two circular doorways (5cm diameter) within the dividing walls of the chambers allowed the test mouse to access each side chamber. Access to the doorways was permitted or restricted using two removable, white polycarbonate partitions. These partitions were inserted into the middle of the apparatus to shield the side chambers prior to the beginning of each trial. A webcam was mounted over the arena to record each testing session. The entire apparatus was thoroughly cleaned with 70% ethanol between each 30-minute trial. The location of the novel inanimate object and stimulus mouse were counterbalanced between trials to remove bias towards one side of the chamber. Black, wire cups (9.7 cm H x 9 cm W) (pen cup mesh-black, Kmart, Australia) were used to encapsulate either the novel subject or object and was held down by a clean, standard water bottle from the animal holding room. The wire cups allowed for auditory, olfactory, and visual but little to no physical contact. The water bottle prevented the test mice from climbing on top of the wire cups. Wire cups containing social scent were stored in separate storage containers away from novel object cups. Further, social cups were stored based on male or female scent to ensure no male-female scent mixing would confound test mice.

*Stimulus mice*

The mice used as novel stimulus subjects were C57Bl/6 wild type mice, aged 6-14 weeks and matched to test mice by sex. Stimulus mice were habituated to the apparatus and wire cup enclosures 1-week prior to the beginning of experimental testing. The habituation phase for stimulus animals consisted of placement underneath the wire cups for 45-60 minutes inside the dimly lit testing room for 2 consecutive days. This ensures stimulus mice will not feel anxious during the test which could influence the behaviour of the test mouse. On test day, stimulus mice were held in a dimly lit holding room, separate to where test mice were held before the commencement of testing to ensure all encounters remained novel.

*Testing day*

During the habituation trial, an empty wire mesh cup was placed in each side chamber, and the mouse was placed in the centre chamber. The partitions were then lifted simultaneously, allowing the mouse to freely explore all chambers for 10-minutes.

During the sociability trial, each side chamber consisted of either a novel inanimate object (neon yellow golf ball used here) or a novel sex-matched stimulus mouse placed under a wire mesh cup. The test mouse remained inside the centre chamber of the arena while the novel object and novel mouse were placed in either side chamber. Then, the partitions were removed to allow the test mouse to freely explore all chambers for 10-minutes.

During the social novelty trial, the novel object was replaced with a novel (unfamiliar) sex-matched mouse while the test mouse was confined to the centre chamber. The novel mouse from the previous trial remained in the same side chamber and was now considered the familiar mouse. The partitions were removed, and the test mouse was allowed to freely explore all chambers for another 10-minutes.

Parameters analysed from this test included time spent engaging with the novel mouse versus the inanimate novel object and time spent with the familiar mouse versus unfamiliar mouse. Sociability is defined as the test mouse engaging more with the novel stimulus mouse as opposed to an inanimate object. Social novelty is defined as the test mouse engaging more with a novel stimulus mouse opposed to a familiar stimulus mouse (Silverman et al., 2010).

**Marble burying**

Marble burying testing is a valuable test for measuring repetitive digging behaviour in rodents (Silverman et al., 2010; Thomas et al., 2009). Since repetitive behaviours like stereotypies and compulsions are common symptoms in several neurodevelopmental disorders (NDDs) (Whitehouse and Lewis, 2015), this test is clinically relevant and widely used in rodent models of NDDs.

To set up the test, the testing room was dimly lit (< 50 lux) and a standard cage (36 cm x 18 cm x 13 cm) was filled 3-4 cm thick with clean corn cob bedding (Sheperd's™ cob, Biological Associates, NSW Australia) and overlaid with 20 black, glass marbles in a 4 x 5 arrangement. The test mouse was placed in the left-hand corner of the testing cage with the head of the mouse facing the marbles, the cage lid was secured, and the mouse was left for a 30-minute testing period. After 30-minutes, the mouse was removed, and a photo was taken of the bedding. The number of marbles buried - defined as more than 2/3rds or more covered by the bedding - was recorded.

**Self-grooming**

Self-grooming is a complex, innate behaviour in rodents that has become valuable for understanding the aetiology of several NDDs. Excessive self-grooming is characterised by unusually long bouts of licking and scratching the entire body (Silverman et al., 2010).

Each test consisted of placing the test mouse in an empty, standard holding cage (36 cm x 18 cm x 13 cm) for a 20-minute test period. A webcam camera was mounted at the level of the cage and the test commenced with the experimenter out of the room. The first 10-minute period was a habituation period and left unscored. The remaining 10-minute period was manually scored by the experimenter who was blinded to the experimental group. The time spent grooming and percentage of time spent grooming were recorded and analysed.

**Integrated neurodevelopmental disorder (NDD) behavioural index**

Integrated behavioural measures from complimentary tests can be z-normalised, reducing the behavioural noise that is a common artifact of behavioural assays (Guilloux et al., 2011; Kraeuter, 2023). A z-normalised value is unitless and indicative of how many standard deviations a subject is above or below the mean of the control population for a particular test. This integration ultimately provides a summary statistic of the subject's emotional state based on both the emotional context of the tests performed and the number of tests integrated. We have used this method previously to characterise the anxiety-like behavioural profile in our dams that produced the F1 offspring used in this study (Bucknor et al., 2024). Here, we took z-normalised values from elevated plus maze, marble burying and self-grooming output to produce an integrated NDD behavioural index. Specifically, we integrated: the percentage of entries and time spent in the open arms of the maze, percentage of marbles buried, and percentage of time spent grooming. These analyses were sex-independent and calculated in reference to either male (♂) or female (♀) HF-/SIS- control groups. The formula used to calculate z-normalised values for each test is below:

$$z = \frac{X - \mu}{\sigma}$$

($X$) = observed output

($\mu$) = group mean

($\sigma$) = population's standard deviation (♂/♀ HF-/SIS- control group)

To integrate multiple behavioural z-scores to produce an integrated NDD behavioural index, the individual z-scores from each mouse were added together and divided by the number of tests included in the analysis:

$$Integrated\ NDD\ behavoiural\ index = \frac{Z_{EPM} + Z_{MB} + Z_{SG} + Z_{NO} + Z_{FAM}}{number\ of\ tests\ (5)}$$

**Tissue collection for scRNA-sequencing and bulk proteomics**

For scRNA-sequencing and bulk proteomic analyses, peripheral blood leukocytes and forebrain brain tissue were analysed from male offspring. Sample numbers for each experiment and corresponding group can be found in **Table S1**.

For peripheral blood and brain tissue collection, mice were first deeply anesthetised with isoflurane (5% induction, 2.5% maintenance) (IsoFlo®, Abbott Laboratories, Botany, NSW, Australia). Up to 1 mL of peripheral whole blood was collected via cardiac puncture into the right ventricle of the heart and placed into a 1.5 mL collection tube containing heparin (10 kU/ml) (10% volume of blood) (Sigma, catalog # H3393) on ice. Then, the mouse was perfused with 20-25 mL of ice cold 1 X PBS and the whole brain was dissected after decapitation. The olfactory bulbs and cerebellar tissue were excised, and the total forebrain was weighed and recorded before tissue homogenisation to generate a single-cell suspension using the Adult Brain Dissociation kit as per manufacturer's instructions (Miltenyi Biotec, catalog# 130-107-677). To isolate leukocytes from whole blood, red blood cell lysis was performed using cold ammonium chloride solution (StemCell Technologies, catalog# 07850) at a volume:volume ratio of 9:1 (i.e., 9mL ammonium chloride and 1mL blood). The solution was incubated at 4°C for 10-minutes, before cells were washed in DPBS (Dulbecco's phosphate-buffered saline: +$Ca^{2+}$, +$Mg^{2+}$, +glucose, +pyruvate; ThermoFisher catalog# 14287080) and centrifuged at 250 x g for 6-minutes at 4°C.

**Table S1.** Sample group information for 'omics' analyses.

| | | | **Brain** | |
|---|---|---|---|---|
| *Group* | *Median Age* | *Sex* | *scRNA seq (n)* | *Proteomics (n)* |
| HF-/SIS-/PIC- | 13-weeks | ♂ | 2 | 3 |
| HF+/SIS+/PIC+ | 14-weeks | ♂ | 2 | 3 |
| | | | **Blood** | |
| HF-/SIS-/PIC- | 13-weeks | ♂ | 2 | 3 |
| HF+/SIS+/PIC+ | 14-weeks | ♂ | 2 | 3 |

**HIVE CLX™ scRNA-sequencing sample capture and library processing**

Once single-cell suspensions were generated for each sample condition (blood/brain; HF+/SIS+/PIC+/ and HF-/SIS-/PIC-), approximately 30,000 cells were loaded directly into a designated HIVE CLX™ collector (HIVE Collector, Honeycomb Biotechnologies, Inc, USA) in 1% FBS and DPBS media. Single-cells settled into picowells of the HIVE collector containing barcoded mRNA-capture beads. Media was removed from the collector and 2 mL of CLX sample wash solution was added, and then 1 mL cell preservation solution. HIVE collectors corresponding to each sample and group condition were stored at -80°C until transferred to AGRF for single-cell NGS library processing and transcriptome recovery. (Further protocol details for sample capture can be obtained here).

Following manufacturer's instructions, HIVE CLX™ collectors were sealed with a semi-permeable membrane, allowing for the addition of strong lysis solution and hybridization solution. After collection, capture beads with transcripts were extracted from the collector by centrifugation. Remaining library preparation steps were performed in a 96-well format. For library size and quality assessment, TapeStation software was used with the High Sensitivity D5000 ScreenTape assay (Agilent Technologies, CA, USA) and final pooled libraries were determined by qPCR. HIVE scRNA-sequencing libraries were sequenced using specific primers contained in the kit using the Illumina® NovaSeq® X Plus sequencing platform. (Further protocol details for HIVE CLX™ scRNAseq transcriptome recovery and library processing can be found here).

#### *scRNA pathway enrichment analysis*

To identify functional pathways from differentially expressed genes across conditions and unique cell types, we performed overrepresentation enrichment (ORA) of gene ontology (GO). ORA GO pathway analysis was performed using the *compareCluster* function from the *clusterProfiler* package. Significant ORA GO pathways (FDR < 0.05) were further simplified to remove redundancy using the *simplify* function in *clusterProfiler.* All ontologies were included for analysis (BP, MF, CC).

### Tissue lysis, protein digestion and peptide tagging for LC-MS/MS analysis

*(a) Brain tissue*

Tissue samples were thawed by incubating at 85°C for 5 minutes for proteome analysis. 100 µL of 10% SDS in water and additional water was added to make up a 400 µL total volume. Samples were incubated again at 85°C for 10 minutes with 10 mM TCEP, sonicated (Branson, Sonifer 150 W, micro tip default setting, 15 s) and then cooled. Samples were incubated at 23°C for 30 minutes with 20 mM iodoacetamide, then precipitated using the chloroform-methanol method (Wessel and Flügge, 1984). Pellets were dried at 37°C for 1h, then 20 µL of 7.8 M Urea/100 mM HEPES pH 8.0/LysC solution was added (3 µg LysC) (Fujifilm, Wako). LysC digestion was for 12h at 28°C. The sample was diluted 8-fold with 100 mM HEPES pH 8.0 and two trypsin digestions were done for 8h at 28°C, each with 3 µg of trypsin (TrypZean, Sigma). The approximate amount of protein was determined by UV absorption at 280 nM (Implen Nanophotometer N60). Tandem mass tag (TMT) labelling was performed using TMT10plex reagents (ThermoFisher Scientific, Cat#90110) on 250 µg of each sample: total negative controls (HF-/SIS-/PIC-): 126, 127N, 127C; total positive samples (HF+/SIS+/PIC+): 128N, 128C, 129N. The combined TMT-labelled sample was cleaned and desalted using a solid phase extraction (SPE) cartridge (Sep-Pak Vac 3cc 200 mg tC18, Waters, Cat#WAT054925). Approximately 150 µg of the combined peptide sample was applied to hydrophilic interaction liquid chromatography separation (HILIC) as described previously (Engholm-Keller et al., 2019). HILIC separation was done using a Vanquish Neo HPLC system with a 250 mm long and 1 mm inside diameter TSK gel Amide-80 column (Tosoh Biosciences). Fractions were collected into a 96-well plate using an FC204 fraction collector (Gilson) at 1-min intervals and monitored by absorbance of UV at 214 nm. Selected fractions were combined into similar amounts of peptide, as indicated by the UV signal, which were dried and reconstituted in 0.1% formic acid for LC-MS/MS analysis.

*(b) Peripheral blood*

The same procedure was carried out as described above with minor changes. Samples were thawed by incubating at 85°C for 5 minutes, then cooled and incubated at 37°C for 30 minutes with 10 units of benzonase. 100 µL of 10% SDS was added and $H_2O$ to make up a 400 µL total volume. Samples were incubated again at 85°C for 10-mins with 10 mM TCEP, sonicated and then cooled. Samples were incubated at 23°C for 30 minutes with 20 mM iodoacetamide, then precipitated using the chloroform-methanol method (Wessel and Flügge, 1984). Pellets were dried at 37°C for 1h, then 10 µL of 7.8 M Urea/100 mM HEPES pH 8.0/LysC solution was added (1 µg LysC). LysC digestion was for 12h at 28°C. The sample was diluted 8-fold with 100 mM HEPES pH 8.0 and two trypsin digestions were done for 8h at 28°C, each with 1 µg of trypsin. The approximate amount of protein was determined by UV absorption at 280 nm. TMT labelling was performed using TMTpro reagents (ThermoFisher Scientific, Cat#A44522) on 23 µg of each sample: total negative controls (HF-/SIS-/PIC-): 126, 127N, 128N; total positive samples (HF+/SIS+/PIC+): 129N, 130N, 131. The combined TMT-labelled sample (138 µg) was cleaned and desalted using a smaller SPE cartridge due to smaller peptide amount (Oasis Prime-HLB 1cc 30 mg, Waters, Cat#186008055). The eluate was applied to HILIC as described above and fractions of peptide were dried and reconstituted in 0.1% formic acid for LC-MS/MS analysis.

**LC-MS/MS analysis**

The LC-MS/MS was performed using a Dionex UltiMate 3000 RSLC nano system and Q Exactive Plus hybrid quadrupole-orbitrap mass spectrometer (ThermoFisher Scientific). Each HILIC fraction was loaded directly onto an in-house 300 mm long 0.075 mm inside diameter column packed with ReproSil Pur C18 AQ 1.9 µm resin (Dr Maisch, Germany). The column was heated to 50°C using a column oven (PRSO-V1, Sonation lab solutions, Germany) integrated with the nano flex ion source with an electrospray operating at 2.3 kV. The S lens radio frequency level was 50 and capillary temperature was 250°C.

*Proteomics of brain and peripheral blood leukocytes*

Samples from brain or peripheral blood were injected in 3.5 µL and loaded onto the column for 17.5-mins at 300 nL/min using 99% buffer A (1% solution of 0.1% formic acid) and 1% buffer B (solution of 0.1% formic acid, 90% acetonitrile). The gradient at 250 nL/min was from 1% buffer B to 7% buffer B in 1 min, to 29% buffer B in 101.5 min, to 36% buffer B in 8 min, to 99% buffer B in 1 min, held at 99% buffer B for 2 min, to 1% buffer B in 1 min and held for 8 min as the flow rate increased to 300 nL/min. MS acquisition was performed for 140 min.

All blood and brain samples and fractions were analysed using data-dependent acquisition LC-MS/MS. MS scans were performed at 70,000 resolution with an AGC target of 1,000,000 and max ion time of 100 ms (m/z 375-1500). MS/MS scans were at 35,000 resolution with an AGC target of 200,000 and max ion time of 100 ms. Parameters included a loop count of 12, 1.1 m/z isolation window, first mass at m/z 120, and collision energy of 31. Singly charged ions and those >8+ were excluded, with a 35 s dynamic exclusion.

**Differential abundance protein and pathway enrichment analysis**

For computational analyses, raw LC-MS/MS data was processed with MaxQuant v1.6.7.0. Variable modifications were oxidation (M), acetyl (protein N-terminus) and deamidation (NQ). Carbamidomethyl (C) was a fixed modification. Digestion was set to trypsin/P with a maximum of 3 missed cleavages. The TMT10plex or TMTpro correction factors were entered (Lot# QB211242 and Lot# TJ268160). Minimum reporter peptide ion fraction was 0.6. The *Mus Musculus* reference proteome with canonical and isoform sequences downloaded Oct 5, 2023, with 55,087 entries and 21,866 genes. The inbuilt contaminants FASTA file was used, and second peptides search was enabled. The peptide spectrum matching, and protein false discovery rates were set at 1%. All modified peptides and counterpart non-modified peptides were excluded from protein quantification. All other settings were default.

*Data cleaning*

The 'proteinGroups.txt' output file from MaxQuant were processed and each protein group must have had at least one unique peptide. Proteins were removed from further analysis if they matched contaminant entries or reverse sequence decoy database in which the protein accession number began with CON_ or REV_ prefixes, respectively. Proteins with one or missing values in any samples were also removed from further analysis. The 'reporter intensity corrected' column was used for further analysis.

*Data normalisation*

UniProt accession was identified for each protein group following rules from Engholm-Keller et al (2019). Multi-mapped proteins were excluded to avoid false enrichment due to alignment errors. Of the accepted proteins, 88.5 % had 1 unique peptide and 11.4 % had more than 2 unique peptides. Samples were first $\log_2$ transformed, then between sample normalisation was performed using the *scaled* normalisation from the *limma* R package (v4.2.3). To remove batch effects from biological samples, the remove unwanted variation *RUV* R package was used (Gagnon-Bartsch et al 2012). RUV relies on having a set of endogenous negative control proteins which have small changes in protein abundances between different cell types or experimental conditions. For this study, a set of 500 empirical negative control proteins with little or no change in protein expression across samples (q-value <0.05) was identified from an initial ANOVA test. The RUVIII method was then used to remove unwanted variations across the samples and four unwanted components (k = 4) were removed by the tool. Differential abundance analysis of proteins was performed using the adjusted abundance matrix.

*Linear modelling*

Differential abundance analysis of proteins was performed for comparing each time point with zero using the *limma* R package. The linar model for comparing each pair of time points was fitted using the *lmFit* function and p-values were calculated using the empirical Bayes method *eBayes* function. Trended and robust analysis were enabled. The Benjamin-Hochberg false discovery rate (FDR) correction was applied to the moderated p-values by calculating q-values (Storey, 2002). Significant differentially expressed proteins (q-value < 0.05) were analysed for GO pathway enrichment analysis using the *enrichGO* function from the *clusterProfiler* R package.
